# Intestinal Paneth cell differentiation relies on asymmetric regulation of Wnt signaling by Daam1/2

**DOI:** 10.1101/2023.01.24.525366

**Authors:** Gabriele Colozza, Heetak Lee, Alessandra Merenda, Szu-Hsien Sam Wu, Andrea Català-Bordes, Tomasz W. Radaszkiewicz, Ingrid Jordens, Ji-Hyun Lee, Aileen-Diane Bamford, Fiona Farnhammer, Teck Yew Low, Madelon M. Maurice, Vítězslav Bryja, Jihoon Kim, Bon-Kyoung Koo

## Abstract

The mammalian intestine is one of the most rapidly self-renewing tissues, driven by actively cycling stem cells residing at the crypt bottom^1,2^. Together with stromal cells, Paneth cells form a major element of the niche microenvironment that provides various growth factors to orchestrate intestinal stem cell homeostasis, such as Wnt3^3^. With 19 family members, different Wnt ligands can selectively activate β-catenin dependent (canonical) or independent (non-canonical) signaling^4,5^. Here, we report that Dishevelled-associated activator of morphogenesis 1 (Daam1) and its paralogue Daam2 asymmetrically regulate canonical and non-canonical Wnt (Wnt/PCP) signaling, and their function is required for Paneth cell progenitor differentiation. We found that Daam1/2 interacts with the Wnt antagonist Rnf43, and Daam1/2 double knockout stimulates canonical Wnt signaling by preventing Rnf43-dependent endo-lysosomal degradation of the ubiquitinated Wnt receptor, Frizzled (Fzd). Moreover, single-cell RNA sequencing analysis revealed that Paneth cell differentiation is impaired by Daam1/2 depletion, as a result of defective Wnt/PCP signaling. Taken together, we identified Daam1/2 as an unexpected hub molecule coordinating both canonical and non-canonical Wnt signaling, the regulation of which is fundamental for specifying an adequate number of Paneth cells while maintaining intestinal stem cell homeostasis.

## Introduction

The intestinal epithelium provides both a large surface for nutrient uptake as well as a physical barrier against harmful agents such as pathogens, and during homeostasis new cells are rapidly generated to maintain its functionality^1,2^. It is organized in a villus-crypt structure consisting of numerous cell types, including intestinal stem cells (ISCs) and Paneth cells (PCs) at the crypt bottom. Previous studies have shown that multiple signaling pathways are involved in the homeostatic turnover and differentiation of the intestinal epithelium, with Wnt and Notch signaling playing a key role in ISC maintenance and differentiation^1^. The potency of Wnt signaling depends mainly on the cell surface levels of its receptor Fzd, which in turn is controlled by Rnf43 and Znrf3 (RZ), transmembrane E3 ligases that ubiquitinate Fzd and promote its degradation through the endo-lysosomal system, thus tightly controlling Wnt signals^6–8^. RZ are in turn regulated negatively by the R-spondin (Rspo) ligand-Lgr4/5 receptor complex^7,9,10^, and positively by phosphorylation of their cytoplasmic tail^11^. In the intestinal epithelium, PCs are the main source of Wnt and Notch ligands^12^. Notch signaling controls cell fate determination of the absorptive and secretory lineages, while canonical Wnt signaling regulates ISC maintenance and proliferation^1^. However, it is still unclear how non-canonical Wnt signaling influences intestinal epithelial homeostasis and differentiation and in which way the non-canonical Wnt signaling is coordinated with canonical Wnt signaling. Here we demonstrate that Daam1 and its paralogue Daam2, members of the diaphanous-related formin (DRF) family of Rho GTPase effectors^13^, function as hub molecules for optimal canonical vs. non-canonical Wnt signaling, to maintain appropriate numbers of ISCs and PCs in the intestinal crypt.

## Results

### Identification of potential Rnf43 interactors

To identify potential functional regulators of Rnf43, we performed mass spectrometry analysis using Rnf43 as bait. From this analysis, we captured a number of potential interactors, listed in Supplementary Table 1. To functionally test the role of these candidates in Rnf43 regulation, we performed a small-scale CRISPR/Cas9 knockout screen on mouse small intestinal organoids grown in standard medium containing Wnt, EGF, Noggin and R-spondin 1 (WENR) (Fig. 1a-f). Guide RNAs were designed to knock out paralogue genes as well, in order to prevent genetic compensation caused by functional redundancy^14,15^ (Fig. 1c). For this purpose, we first established a mouse small intestinal organoid line expressing Rnf43-IRES-mCherry (Dox-Rnf43 SI organoid) under the control of a doxycycline responsive promoter (Fig. 1a, b). As expected, in the presence of doxycycline, these organoids rapidly died as a consequence of Wnt blockade by overexpressed Rnf43 (Fig. 1b, d). However, genetic depletion of the putative Rnf43 interactors could prevent organoid death by maintaining high levels of canonical Wnt signaling, similar to the knockout (KO) of other negative regulators of Wnt/β-catenin, such as Axin1/2 (Fig. 1b, e and f). Using this screening platform, we identified Daam1 and Daam2 as functional interactors of Rnf43 (Fig. 1f, h). Depletion of Daam1/2 (D1/2 DKO) in Dox-Rnf43 organoids could rescue organoid survival as efficiently as Axin1/2 KO (Axin DKO), despite the presence of Rnf43 overexpression (Fig. 1f). To test whether Daam1/2 is downstream of the RZ axis (Lgr4/5-Rspo-RZ), we used withdrawal assays for either R-spondin 1 (EN medium) or both Wnt and Rspo (EN + the Wnt inhibitor IWP2) (Fig. 1g). As expected, CRISPR-generated Axin DKO organoids survived in both conditions, whereas RZ DKO organoids did not survive without Wnt (Fig. 1h and Extended Data Fig. 1). Interestingly, in contrast to *wild type* (WT) organoids, D1/2 DKO organoids did survive in Rspo deficient medium (EN), but they also died in the Wnt and Rspo deficient condition (EN+IWP2) (Fig. 1h). These findings suggest that unlike Axin and Apc, which are components of the destruction complex downstream of Wnt and Fzd, Daam1/2 acts downstream of the RZ axis, disruption of which renders organoids independent of Rspo but still dependent on Wnt stimulation for optimal canonical Wnt signaling.

**Fig. 1.**
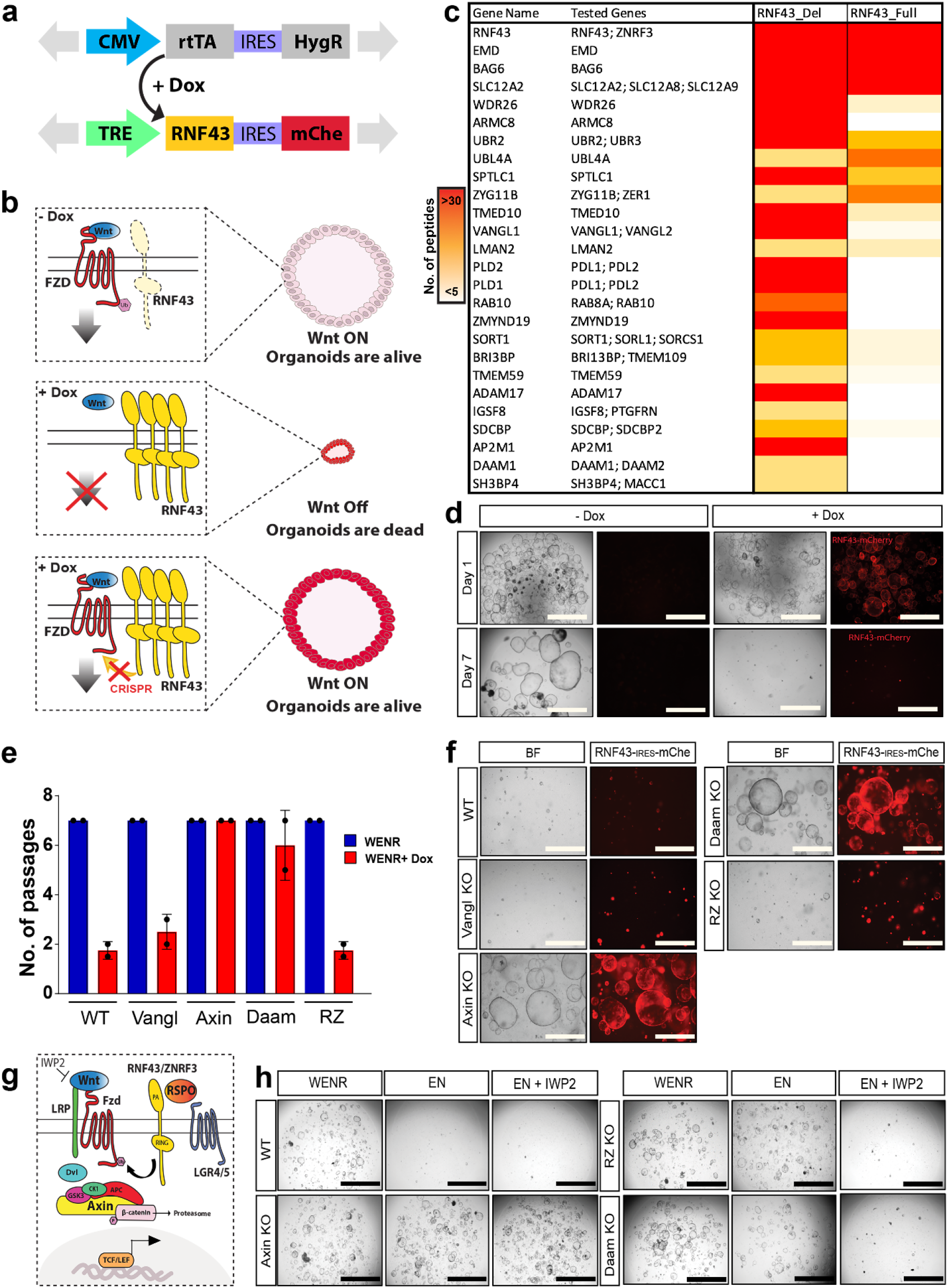
Functional CRISPR screening of Rnf43-interacting proteins. **a,** Schematic of the DNA constructs used to generate small intestinal organoids expressing doxycycline-inducible *Rnf43-IRES-mCherry*. mCherry is used as a proxy for Rnf43 expression. **b,** Schematic showing the CRISPR-based screening of Rnf43-expressing organoids. Doxycyclin addition turns on Rnf43 expression and organoids die, unless downstream components necessary for Rnf43 activity are eliminated via CRISPR/Cas9 KO. **c,** List of Rnf43-interacting proteins identified via mass spectrometry. Abundance of each protein is represented as the number of peptides detected, according to the color scale on the left. **d,** Organoid assay showing robust Rnf43 expression upon doxycyclin treatment. Note that at day 7 all organoids expressing Rnf43 (here visualized through mCherry fluorescence) are dead. **e,** Bar plot quantification of organoid survival after indicated passages. Only Axin and Daam CRISPR-KO organoids grow in the presence of Rnf43 overexpression. **f,** Survival assay of WT and indicated CRISPR KO mutant organoids after Rnf43 induction. **g,** Schematic illustrating the growth factor withdrawal assay used in h, to determine Rnf43-interactor epistasis in the Wnt pathway. **h,** Growth factor withdrawal assay on WT and indicated CRISPR mutant organoids. All scale bars represent 1000 μm.

### Daam1/2 is essential for Rnf43-mediated Fzd endocytosis

Next, we sought to understand how Daam1/2 regulates canonical Wnt signaling in cooperation with Rnf43. Rnf43, as well as its paralogue Znrf3, is a well-known E3 ubiquitin ligase that targets Fzd for ubiquitination-mediated endo-lysosomal degradation^6,7^. Hence, we decided to monitor cell surface levels of Fzd in the absence or presence of Daam1/2. To this aim, we knocked out *Daam1/2* in HEK293T cells (D1/2 DKO HEK293T cells) via CRISPR/Cas9 (Extended Data Fig. 2a, b) and performed receptor internalization assays using a SNAP-tagged Fzd5 construct (Fig. 2a)^6^. In WT HEK293T cells, SNAP-Fzd5 was rapidly internalized and decreased from the plasma membrane when co-expressed with Rnf43. In contrast, D1/2 DKO HEK293T cells maintained higher levels of cell surface Fzd5. We also assessed the endogenous levels of cell surface Fzd receptors by flow cytometry analysis using a pan-Fzd antibody^16^. Even in the presence of Rnf43, D1/2 DKO HEK293T cells showed increased surface levels of Fzd compared to WT cells, where Fzd levels were downregulated upon Rnf43 overexpression (Fig. 2b). Moreover, using a cell-surface protein biotinylation assay, we found that surface levels of the Wnt co-receptor Lrp6 were also retained in the D1/2 DKO HEK293T cells in the presence of Rnf43 (Fig. 2c, lane 5).

**Fig. 2.**
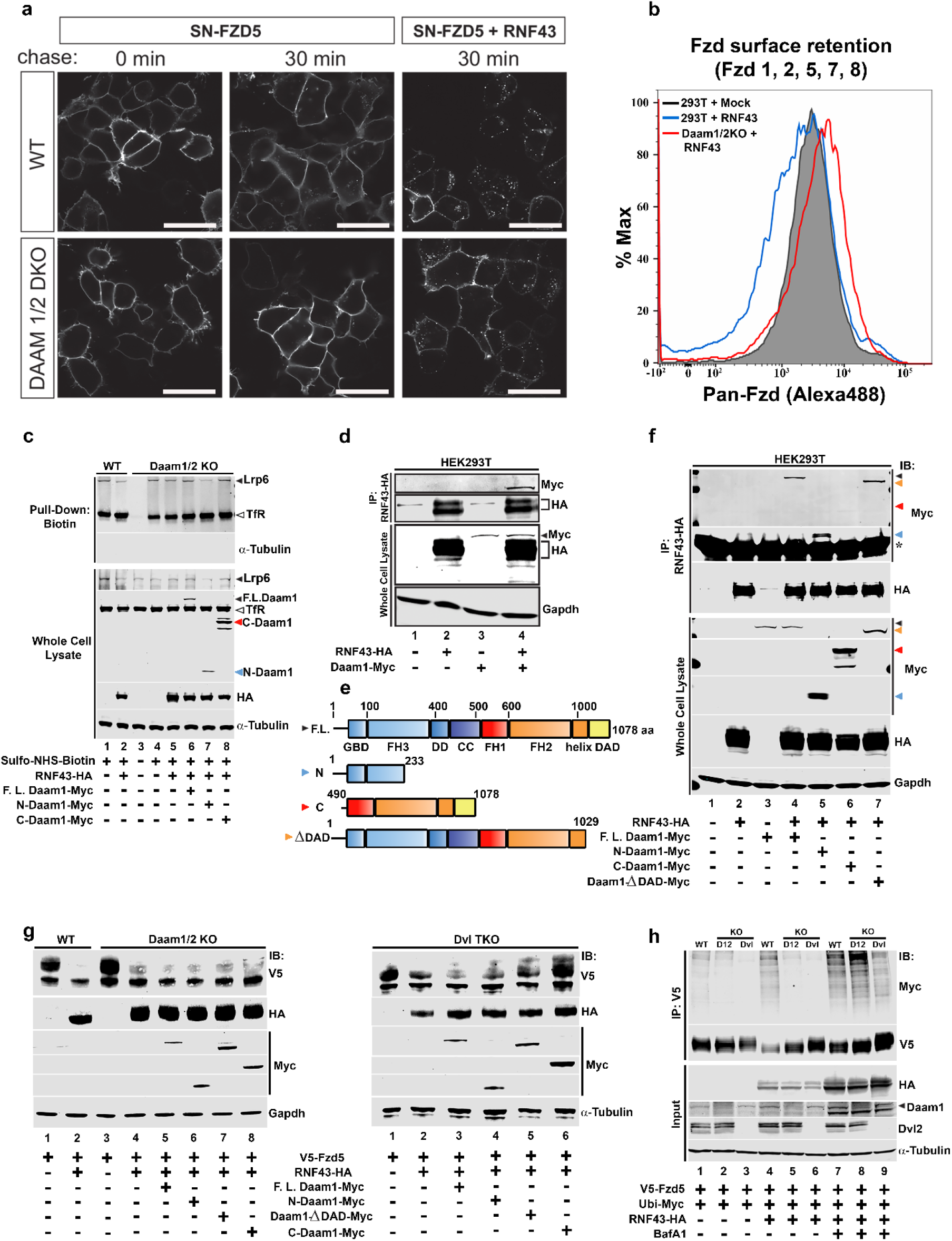
Daam is required for Rnf43-dependent Frizzled endocytosis. **a,** Subcellular localization of SNAP–Fzd5 in WT or Daam1/2 DKO HEK293T cells co-transfected with control empty vector or Rnf43. Surface SNAP–Fzd5 was labeled with SNAP–Alexa549 for 15 min and chased for 30 min. Scale bars represent 20 μm. **b,** FACS analysis of plasma membrane levels of Fzd receptors in WT and D1/2 DKO HEK293T cells, with or without Rnf43 overexpression. **c,** Western blot of representative cell surface protein biotinylation and pull-down assay on HEK293T cells transfected with indicated constructs, showing that Daam1/2 KO also prevents Rnf43-dependent internalization of the Wnt co-receptor Lrp6. Transferrin receptor (TfR) was used as a negative control. **d,** IP assay used to show interaction between HA-tagged Rnf43 and Myc-tagged Daam1. **e,** Schematic of the Daam1 full length and deletion constructs used in this study to map the Rnf43-interacting domain of Daam1. Daam1 architecture domain and relative amino acid position are indicated. GBD, Rho <u>G</u>TPase <u>b</u>inding <u>d</u>omain; FH3, <u>F</u>ormin <u>h</u>omology <u>3</u> domain; DD, <u>d</u>imerization <u>d</u>omain; CC, <u>c</u>oiled <u>c</u>oil domain; FH1, <u>F</u>ormin <u>h</u>omology <u>1</u> domain; FH2, <u>F</u>ormin <u>h</u>omology <u>2</u> domain; Helix, an amphipathic helix involved in interaction with FH3 domain; DAD, <u>D</u>iaphanous <u>a</u>utoregulatory <u>d</u>omain. **f,** IP experiment showing that the N-terminal domain of Daam1 is required for Rnf43 interaction. Colored arrowheads correspond to the different Daam1 constructs, as shown in panel e, and indicate their migration position on the blot. **g,** Western blots showing Frizzled degradation by Rnf43 in WT, D1/2 DKO and Dvl TKO HEK293T cells. **h,** Western blot showing ubiquitin levels of Fzd5 in WT, D1/2 and Dvl mutant cells. α-tubulin was used as a loading control in c, g and h. Gapdh was used as a loading control in d, f and g.

To precisely understand the role of Daam1/2 in Rnf43-mediated Fzd endocytosis, we investigated Rnf43-Daam1/2 interactions and Fzd ubiquitination. Firstly, immunoprecipitation assay using overexpressed full-length (F.L.) tagged Daam1 and Rnf43 showed that Rnf43 could efficiently pull down myc-tagged Daam1 (Fig. 2d), corroborating our mass spectrometry data (Fig. 1c). Then, to identify which domain of Daam1 is required for Rnf43 interaction, we overexpressed full-length Daam1, previously described^17^ N-term and C-term truncated constructs, as well as a Daam1 mutant lacking the C-terminal Dvl-interacting^18^ diaphanous autoregulatory domain (DAD) domain (Fig. 2e), together with Rnf43 in HEK293T cells. We were able to precipitate Rnf43 together with full-length (F.L.), N-term, and ΔDAD mutant Daam1 but not with C-term Daam1 (Fig. 2f), indicating that the Daam1 N-terminal domain is necessary and sufficient for its interaction with Rnf43. Notably, overexpression of the N-terminal domain, but not the C-terminal domain, could also rescue Rnf43-mediated clearance of Lrp6 from the plasma membrane (Fig. 2c, lanes 7 and 8). Only partial effect was observed with F.L. Daam1 (Fig. 2c, lane 6), in line with previous studies reporting that Daam proteins, like other formins, stay in a closed, auto-inhibited conformation^13,17,18^.

Next, we checked the effect of D1/2 DKO on Rnf43-mediated Fzd ubiquitination and degradation. As Dishevelled1/2/3 (Dvl1/2/3) are known to mediate the interaction between Fzd and Rnf43^19^, we first compared the degradation of Fzd in D1/2 DKO and Dvl triple KO (TKO) HEK293T cells. In the presence of Rnf43, the mature form of Fzd was specifically downregulated in WT HEK293T cells (Fig. 2g). In D1/2 DKO HEK293T cells, the levels of mature Fzd were reduced but not completely suppressed (Fig. 2g). The same was observed in the Dvl TKO HEK293T cells (Fig. 2g), as expected since Dvl bridges the interaction between Rnf43 and Fzd^19^. This indicates that Daam1/2, like Dvl, is necessary for Rnf43-mediated degradation of Fzd.

However, differences were observed between D1/2 DKO HEK293T and Dvl TKO HEK293T in the levels of Fzd ubiquitination: in Dvl TKO cells, Fzd ubiquitination was clearly reduced (Fig. 2h), due to the lack of bridging between Rnf43 and Fzd^19^. In contrast, D1/2 DKO showed elevated levels of Fzd ubiquitination, even when compared to that of WT HEK293T cells (Fig. 2h), suggesting that the Fzd ubiquitination is still intact in D1/2 DKO HEK293T cells. We speculate that Daam1/2 is necessary for the internalization and subsequent vesicle sorting of ubiquitinated Fzd receptors, whereas Dvl is important for Rnf43-Fzd interaction and ubiquitination of Fzd. Taken together, we propose the following stepwise mechanism of ubiquitination-mediated Fzd endo-lysosomal degradation: first, Dvl promotes the interaction between Rnf43 and Fzd, leading to Fzd ubiquitination, after which Daam1/2 promotes the internalization and subsequent sorting of ubiquitinated Fzd.

### Daam1/2 deletion confers Rspo independence but fails to phenocopy Rnf43/Znrf3 deletion

To assess the consequences of Daam1/2 deletion *in vivo* in the mouse intestine, we generated a Daam1/2 double conditional knockout mouse model (D1/2 cDKO), specifically for the small intestine by using the Villin-CreERT2 (Vil-CreER) transgenic system (Extended Data Fig. 2c-e). We injected tamoxifen in 6 – 8 weeks old, age-matched WT, D1/2 cDKO and RZ cDKO mice to ablate Daam1/2 and RZ in the small intestinal epithelium, respectively. Two weeks after tamoxifen injection, small intestine sections were examined by Ki67 and lysozyme (Lyz) immunostaining to detect proliferative progenitor cells and Paneth cells, respectively. As previously reported, RZ cDKO small intestinal epithelium showed clear hyperplasia with increased numbers of Ki67+ proliferative cells and Lyz+ Paneth cells (Fig. 3a)^6^. However, D1/2 cDKO epithelium only showed a mild increase of both cell types (Fig. 3a), and there was no clear sign of hyperplasia in any of the examined sections.

**Fig. 3.**
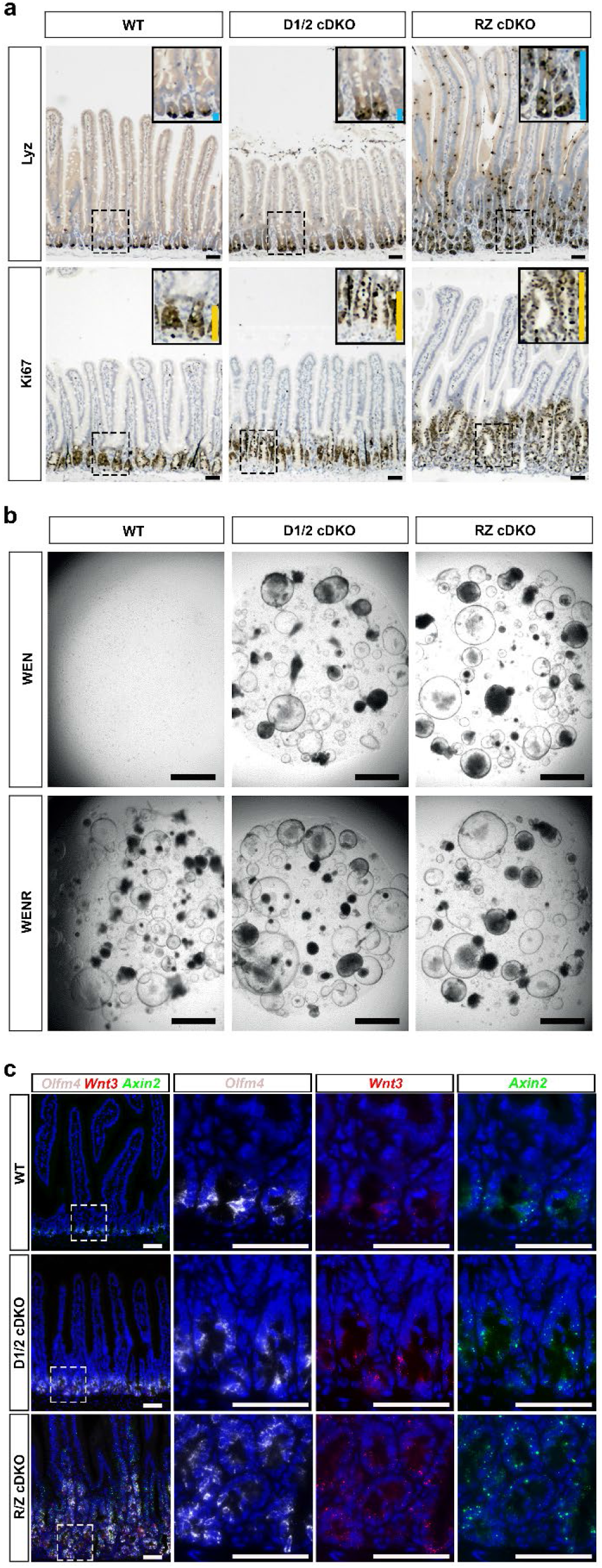
Daam1/2 cDKO mice show a milder phenotype than RZ cDKO, despite maintaining Rspo1 independence. **a,** Small intestine histological sections from WT, D1/2 and RZ conditional double knockout mice stained for lysozyme (Lyz), a Paneth cell marker, and Ki67, used as a proliferation marker. Insets show magnification of dash boxed areas. The extent of Lyz-positive Paneth zones and Ki67-positive proliferative zones are indicated by blue and yellow side bars, respectively. **b,** Organoids derived from WT, D1/2 and RZ cDKO mice, showing that D1/2 and RZ mutant organoids can survive in the absence of R-Spondin, unlike WT organoids. **c,** RNA-scope *in situ* hybridization analysis with probes targeting *Olfm4* (stem cell marker), *Wnt3* (Paneth cell marker) and *Axin2* (canonical Wnt target). White dashed boxed areas shown in the left “merge” panels are enlarged and shown on the right as single probe stainings. Scale bars represent 50 μm in a and c, and 1000 μm in b.

We next tested whether cells from the newly generated D1/2 cDKO mouse showed Rspo independence, as we observed in Daam1/2 CRISPR mutants. To this end, we generated organoids from the small intestine of WT, D12 cDKO, and RZ cDKO mice after tamoxifen-induced deletion of Daam1/2 and RZ, respectively. All isolated organoids with different genotypes grew well in WENR medium. When grown in WEN, only D1/2 and RZ cDKO organoids survived (Fig. 3b), confirming that D1/2 cDKO organoids acquired Rspo independence, as observed in the original D1/2 DKO CRISPR mutant organoids (Fig. 1f). Our organoid data also rules out any other defect in intestinal stem cell maintenance, as D1/2 cDKO organoids could be maintained *in vitro* for multiple passages, as with the other lines. This was also supported by *in situ* hybridization analysis *of* intestinal tissue. *Olfm4* and *Axin2* expression patterns clearly confirmed the presence of intestinal stem cells and canonical Wnt signaling activity, respectively, in all three genotypes (Fig. 3c). Both D1/2 cDKO and RZ cDKO showed elevated *Axin2* signals, again confirming higher canonical Wnt activity (Fig. 3c). However, we noted that *Wnt3+* Paneth cells were not increased in D1/2 cDKO, while Paneth cell hyperplasia was evident in RZ cDKO (Fig. 3c) similar to Lyz immunostaining (Fig. 3a). We conclude that D1/2 cDKO renders intestinal stem cells more sensitive to Wnt ligand stimulation by compromising RZ-mediated negative feedback regulation. However, unlike RZ cDKO, D12 cDKO did not show a concomitant increase in the number of Paneth cells.

### Asymmetric role of Daam1/2 in non-canonical Wnt pathway for differentiation of Paneth cells

Daam1/2 is a well-characterized effector of the non-canonical Wnt/PCP signaling pathway, working downstream of another Wnt signaling regulator, Dvl^17,18^. In particular, Daam proteins were shown to mediate Wnt/PCP-dependent reorganization of the actin cytoskeleton by activating Rho GTPases, a process required for gastrulation movements during vertebrate development^17,18^. Recently, it has also been reported that Paneth cell differentiation involves activation of the Wnt/PCP pathway^20^. This prompted us to test whether the difference in Paneth cell number between D1/2 and RZ cDKO small intestines was caused by the lack of Wnt/PCP signaling.

Hence, we monitored activation of the Wnt/PCP signaling pathway upon Wnt5a stimulation, a well-known non-canonical Wnt ligand, in WT and D1/2 DKO HEK293T cells. Dvl phosphorylation was intact in both WT and D1/2 DKO HEK293T cell lines, indicating proper Wnt/PCP activation upstream of Daam1/2 upon Wnt5a treatment (Fig. 4a and Extended Data Fig. 3a). However, downstream signaling events were compromised as we could not observe any Wnt5a-dependent elevation of active Rho protein level in D1/2 DKO HEK293T cells (Fig. 4a), in agreement with previous observations^17^. In addition, molecular imaging of F-actin dynamics using Lifeact-GFP or phalloidin also showed that D1/2 DKO HEK293T cells lack Wnt5a stimulation-dependent cytoskeleton rearrangement and filopodia formation (Fig. 4b and Extended Data Fig. 3b), a hallmark of Wnt/PCP defect^21–23^. Altogether, these data show that D1/2 KO cells show defects in the Wnt/PCP pathway, in agreement with published work. In contrast, as shown above, with the lack of negative feedback regulation by Rnf43, canonical Wnt signaling activity was enhanced in the D1/2 DKO HEK293T cells as they showed a higher level of unphosphorylated active β-catenin level at a steady state and upon Wnt3a stimulation (Fig. 4c). We also confirmed elevated expression of the canonical Wnt signaling target genes, *Sp5* and *Axin2*, by RT-qPCR (Fig. 4d). To test whether our findings on Wnt/β-catenin modulation by Daam1/2 could be extended to other systems, we resorted to the frog model *Xenopus laevis*. During *Xenopus* development, a Wnt/β-catenin gradient is instrumental in determining the antero-posterior (AP) body axis, where Wnt activity is lower in the anterior and higher in the posterior^24^. Because of this, overexpression of Wnt inhibitors in frog embryos causes enlargement of the head and anterior structures (anteriorization)^25,26^, while Wnt activation induces head loss (posteriorization)^27^. To test if D1/2 knock-down could mimic a Wnt activation phenotype, we performed a D1/2 morpholino-mediated knock-down experiment (Extended Data Fig. 4a-c). When injected into the dorsal blastomeres of 4-cell stage *Xenopus* embryos (Extended Data Fig. 4d), Daam1/2 specific morpholinos synergistically reduced head development compared to WT controls (Extended Data Fig. 4e-g), phenocopying canonical Wnt overactivation (Extended Data Fig. 4h).

**Fig. 4.**
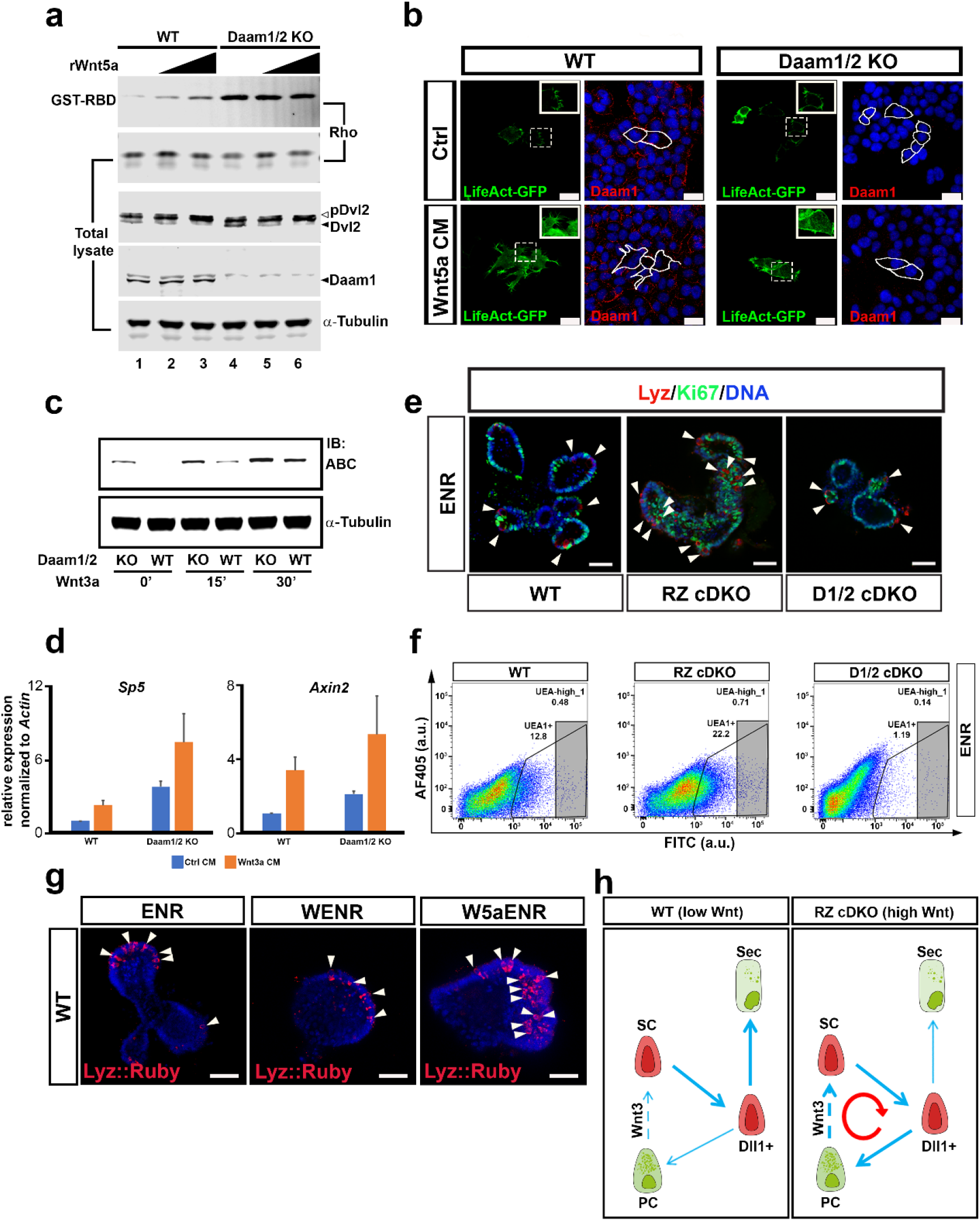
Daam knockout impairs non-canonical Wnt while enhancing canonical Wnt signaling. **a,** Active Rho pull-down assay. WT and D1/2 KO cells were treated with recombinant Wnt5a (0, 200 or 400 ng/ml) for 30 minutes at 37 °C prior to Western blot analysis. White and black arrowheads point to phosphorylated and unphosphorylated Dvl2, respectively, which served to monitor for Wnt5a activity. Daam1 immunoblot was used to confirm its absence in D1/2 KO cells. **b,** WT and D1/2 KO HEK293T cells transfected with LifeAct-GFP were treated with Wnt5a CM for 2 hours at 37 °C, before immunofluorescence analysis. Insets show magnification of the dash boxed area. White lines indicate transfected cell location among non-transfected cells. Scale bars represent 20 μm. **c,** Western blot comparing levels of active β-catenin in WT and D1/2 KO HEK293T cells. Cells were treated with Wnt3a CM for the indicated time, prior to analysis. α-tubulin was used as a loading control in a and c. **d,** RT-qPCR analysis of canonical Wnt target genes *Sp5* and *Axin2* on WT and D1/2 KO cells treated overnight with Wnt3a CM. Expression levels are normalized to *Actin* mRNA. Error bars represent standard deviation across three biological replicates. **e,** Organoids derived from WT, RZ and D1/2 cDKO mice, cultured in ENR medium and stained for lysozyme and Ki67. Arrowheads point at Paneth cells (Lyz^+^, in red). **f,** FACS analysis of organoids from indicated mouse genotypes, stained with fluorescein isothiocyanate (FITC)-labeled *Ulex europaeus* agglutinin 1 (UEA-1). **g,** Confocal imaging of lysozyme::Ruby WT SI organoid reporter line, expressing Ruby RFP in Paneth cells. Organoids were cultured in ENR, WENR on Wnt5a-containing ENR. Scale bars in e and g represent 50 μm **h,** Schematic showing Paneth cell-intestinal stem cell double positive feedback in normal homeostatic and Wnt high conditions (such as in RZ cDKO intestine).

Since Daam1/2 KO impairs Wnt/PCP signaling (Fig. 4a, b), we checked the consequence of the Wnt/PCP defect on Paneth cell differentiation using mouse small intestinal organoids. Lyz immunostaining (Fig. 4e) and UEA1 flow cytometry analysis (Fig. 4f) showed that the number of Paneth cells was dramatically decreased in D1/2 cDKO organoids, as compared to WT and RZ cDKO organoids. Consistent with *in vivo* data^6^ (Fig. 3a), RZ cDKO organoids showed typical Paneth cell hyperplasia (Fig. 4e, f). Furthermore, WT intestinal organoids expressing a fluorescent reporter under regulation of the lysozyme promoter showed a higher number of Paneth cells when treated with Wnt5a, suggesting that activation of non-canonical Wnt signaling is sufficient to stimulate Paneth cell differentiation (Fig. 4g). To further confirm the role of Wnt/PCP signaling in Paneth cell differentiation, we performed single-cell RNA sequencing analysis of WT, RZ cDKO, and D1/2 cDKO organoids that were cultured in WENR conditions. In total, we analyzed 8,000 (WT), 13,000 (RZ cDKO), and 9,000 (D1/2 cDKO) cells, and identified 5 main clusters in UMAP (Fig. 5a and Extended Data Fig. 5a-c). These clusters consisted of revival stem cells, *Lgr5+* stem cells, proliferating cells, enterocytes, and mature secretory lineage cells. Due to the use of high canonical Wnt stimulation in organoid culture medium, progenitor cell populations were highly represented; this is a favorable condition that allowed us to gain insights into the initial commitment to different cell lineages of the progenitor cell populations. The number of Lgr5+ cells was comparable among all three genotypes with a small decrease observed in D1/2 cDKO organoids, which was compensated by a concomitant increase in the fraction of revival stem cells (Fig. 5b). In D1/2 cDKO organoids, the most significantly affected populations were the mature secretory cell types (Fig. 5b). Likewise, *Neurog3+* and *Tff3+* secretory lineage progenitor cells (Fig. 5c) were missing in D1/2 cDKO organoids. Interestingly, *Dll1+* early secretory progenitors were still present in D1/2 cDKO organoids (Fig. 5c), suggesting that the first binary commitment to the secretory lineage by Notch signaling is not affected by Daam1/2 KO. We also observed decreased levels of *Clca3b* and *Cfap126 (Flattop*, a known Wnt/PCP target gene^20,28^) (Fig. 5c, d), as well as of the Paneth cell markers *Defa24* and *Defa17* (Fig. 5c). Most interestingly, the number of *Dclk1+* tuft cells was significantly increased in the D1/2 cDKO organoids (Fig. 5c), probably as a result of the block towards other secretory lineages among committed Dll1+ cells. These results were also confirmed through pseudobulk RNA-seq analysis, as shown in Extended Data Fig. 6a-g. These data provide the first genetic evidence that Wnt/PCP is involved in secretory lineage specification, particularly Paneth cell differentiation, as predicted by previous work^20^. In agreement with the data presented here and from others^20^, we propose the following stepwise binary fate decision model for intestinal cell specification: first Notch signaling directs the choice between absorptive and secretory (*Dll1+*) lineages; then Wnt/PCP signaling regulates differentiation between tuft and other secretory lineages (Fig. 5e).

**Fig. 5.**
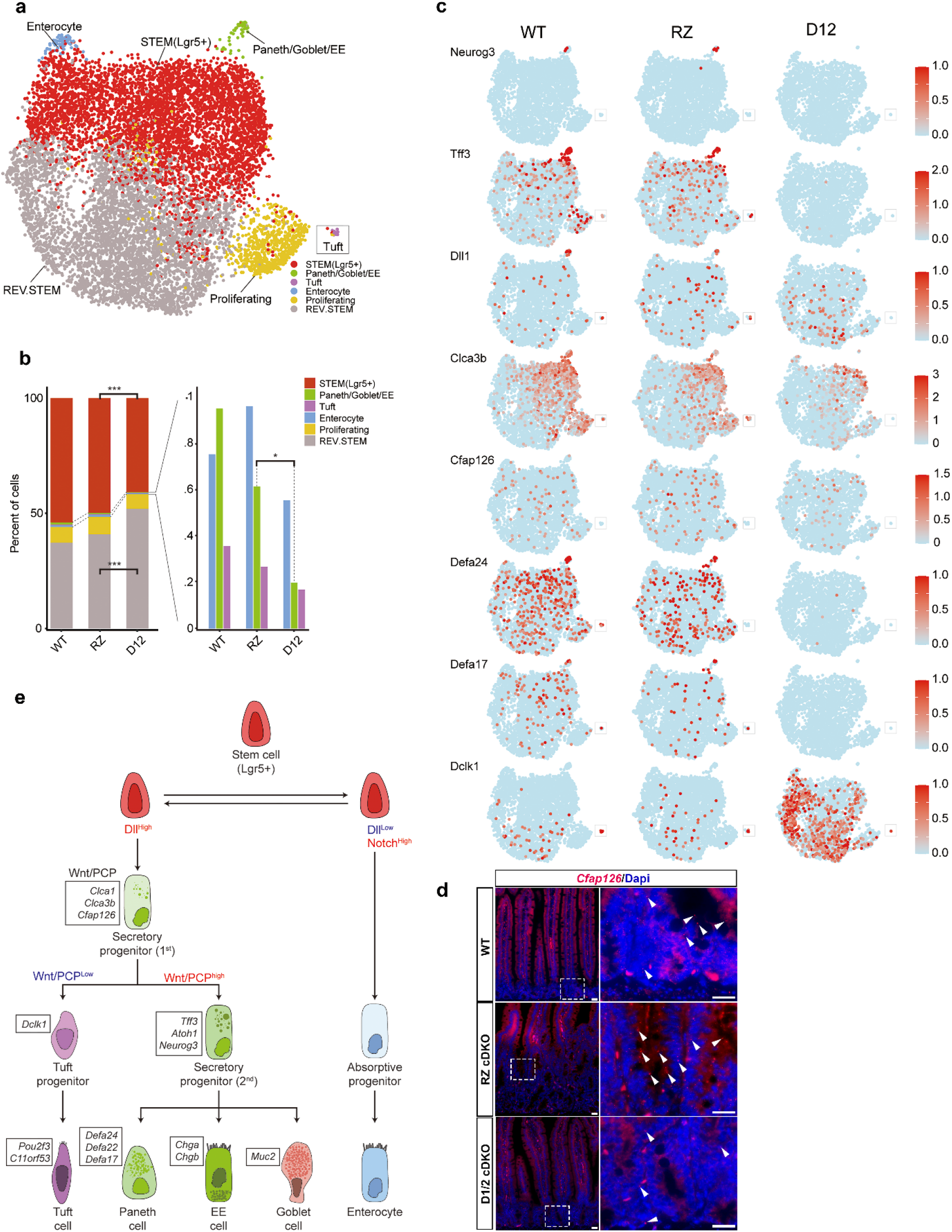
scRNA-seq analysis for WT, RZ and D1/2 cDKO organoids. **a,** Integrated UMAP cluster map including WT, RZ, and D1/2 samples. **b,** Cell-type composition of WT, RZ, and D12 organoids. p-values were calculated by Fisher’s exact tests. Corrected p-values were described as *P < 0.01, ***P < 0.001. **c,** Expression pattern of selected cell type markers on UMAP clusters from individual samples. Red color indicates maximum expression level while blue color indicates minimum expression level for each gene. **d,** *In situ* hybridization using an RNA-Scope probe specific for *Cfap126* (*flattop*) in the small intestinal crypts of WT, RZ and D1/2 cDKO mice. Right panels are enlargement of areas included in the dash boxes on the left. White arrowheads indicate the fluorescent signal from single mRNA transcripts (red puncta). Note the decrease in *Cfap126* expression in D1/2 KO mice. Scale bars represent 20 μm. **e,** Cellular flow chart showing the stepwise commitment from Lgr5+ stem cells. Dll^high^ progenitors can generate secretory cells including tuft, Paneth, enteroendocrine (EE) and goblet cells, while Dll^low^ will only generate enterocytes. Among Dll^high^ progenitors, only cells with active Wnt/PCP signaling can mature into Paneth, enteroendocrine and goblet cells.

Finally, our analysis of the differentiation pattern of D1/2 cDKO intestinal epithelial cells provides a clear explanation for why we did not observe a similar Paneth cell hyperplasia as observed in the RZ cDKO intestine (Fig 3a). We previously showed that introducing Math1 or Wnt3 cKO, which prevents Paneth cell formation, into an RZ cDKO genetic background produced a strongly alleviated phenotype, despite maintaining Rspo independence and Wnt hypersensitivity^29^. D1/2 cDKO leads to the same alleviated phenotype by the unexpected combination of Rspo independence and Paneth cell defects, caused by the asymmetric regulation of canonical (up) and non-canonical (down) Wnt signaling by Daam1/2 KO.

## Discussion

Using cell cultures, organoids and mouse models, we have demonstrated that Daam1 and Daam2 are novel downstream regulators of RZ in the small intestine, which mediate endocytosis of ubiquitinated Fzd. A similar endocytic function for Daam1 has been previously reported and shown to be important for ephrinB signaling in the zebrafish notochord^30^. Of note, interaction with ephrinB occurred via the N-terminal domain of Daam^30^, analogous to the Rnf43-Daam1 interaction uncovered here. Similarly, Daam was shown to interact through its N-terminal domain with β-arrestin2, a protein regulating endocytosis and Wnt/PCP signaling during *Xenopus* convergent extension^31^. We thus speculate that our work uncovers a more general role for Daam proteins in cell signaling regulation via endocytosis, a fundamental process in Wnt signaling^5,32^, perhaps shared by other members of the Formin superfamily^33^.

We have also confirmed that Daam1/2 plays an important role in the non-canonical Wnt pathway as a downstream effector of Dvl. Hence, loss of Daam1/2 enhances canonical Wnt signaling, while reducing Wnt/PCP signaling. In contrast, RZ negatively regulate both canonical and non-canonical Wnt, by targeting canonical Wnt receptors Fzd and Lrp6^6,7^, as well as non-canonical Wnt components Ror1/2 and Vangl1/2^34^ for degradation, such that its loss causes enhanced signaling in both pathways^35,36^. Due to this difference, RZ cDKO intestine showed hyperplasia with over-production of Paneth cells, whereas D1/2 cDKO intestine displayed an alleviated phenotype, characterized by a lack of efficient Paneth cell specification. Our data unveil an unexpected additional binary fate decision step regulated by Wnt/PCP signaling, after the well-known primary fate decision between absorptive and secretory lineages regulated by Notch signaling^1^. This Wnt/PCP-regulated fate determination step seems to govern secretory cell differentiation between tuft cells and other secretory cell types, including Paneth cells (Fig. 5e). In D1/2 cDKO intestine and organoids, secretory lineage differentiation is heavily compromised, particularly at the expense of Paneth cells, while increasing the production of tuft cells, which are usually the rarest population in the gut epithelium.

Paneth cells represent a key epithelial cell population that provides neighboring stem cells with essential niche factors, especially Wnt3. Such epithelial Wnt-producing cells can be an important pro-tumorigenic niche for tumorigenic stem cells that are still dependent on Wnt ligands for their maintenance and growth (e.g., RZ mutants or Rspo-fusion bearing mutants). Paneth cell hyperplasia in RZ tumors is known to create a positive feedback loop that sustains Wnt-addicted tumor cells via the massive production of Wnt-secreting cells^29^ (Fig. 4h). For this reason, porcupine inhibitors have received particular attention as they can be used to inhibit Wnt secretion from the tumorigenic niche^29^. Our data suggest another vulnerable point, since inhibiting non-canonical Wnt signaling can have a similar effect by altering the secretory lineage specification. In conclusion, our study not only provides genetic evidence for a role of Wnt/PCP signaling in intestinal secretory lineage specification, but also opens up new therapeutic avenues to treat Wnt-addicted cancers.

## Author contribution

G.C., A.M., J.K. and B.K. conceived and designed the study. G.C., A.M., S.W., A.C.B., T.W.R., I.J., J.H.L., A.D.B., F.F., T.Y.L. and J.K. performed the experiments. G.C., A.M., S.W., T.W.R., I.J., T.Y.L., M.M.M., V.B., J.K. and B.K. analyzed the results and H.L. analyzed the scRNA-seq data. T.W.R. and V.B. provided material required for this study. G.C., H.L., J.K. and B.K. wrote the manuscript. All authors read and provided comments on the manuscript.

## Acknowledgements

We thank members of the Koo, Elling, and Urban labs for valuable discussions and critical comments, Dr. Rike Zietlow for reading and editing the manuscript, VBC core facilities (especially the Histology Facility, which performed RNA-Scope stainings, BioOptics, Molecular Biology and the animal caretakers), and the lab of Dr. Elly Tanaka for providing the frog embryos. Single-cell RNA-seq was performed by the Next Generation Sequencing Facility at Vienna BioCenter Core Facilities (VBCF), member of the Vienna BioCenter (VBC), Austria. Work in the Koo lab is supported by core funding from the Institute of Molecular Biotechnology (IMBA) of the Austrian Academy of Sciences; ERC starting grant, Troy Stem cells, 639050; Interpark Bio-Convergence Center Grant Program. G.C. is supported by the Austrian Science Fund (FWF), Lise Meitner Program M 2976. S.W. is supported by DOC Fellowship of the Austrian Academy of Sciences. Research in V.B. lab is funded by the Czech Science Foundation (GX19-28347X). This work is dedicated to the memory of Maurizio Colozza (1950-2021).

## Methods

### Mouse husbandry and generation of conditional knockouts

Daam1tm1a(EUCOMM)Hmgu and Daam2tm1a(KOMP)Wtsi chimeras were generated by blastocyst microinjection. Chimeras were crossed with the Villin-CreERT2 (Vil-CreERT2) mouse line to generate the conditional Vil-CreERT2-Daam1fl/fl, Vil-CreERT2-Daam2fl/fl, and Vil-CreERT2-Daam1fl/fl-Daam2fl/fl lines. The Vil-CreERT2-Rnf43fl/fl-Znrf3fl/fl mouse line was included as a positive control. To induce Cre recombinase, 2 mg of tamoxifen in corn oil per 20 g of body weight or corn oil alone for negative control animals were injected at age 8 – 12 weeks. Both male and female mice were included in the experiments. Small intestine crypts were isolated, and organoids were established for further *in vitro* experiments, as described below. All mice were sacrificed two weeks after Cre induction. Standard light/dark cycle, temperature, and humidity parameters were used by the mouse facility to maintain all mouse lines. All animal experiments adhered to the guidelines of the Austrian Animal Care and Use Committee.

### Small intestine organoid establishment and maintenance

Small intestinal crypt isolation and organoid establishment were reported previously ^6,29^. Briefly, mouse small intestinal crypts were isolated by applying Gentle Cell Dissociation Reagent from STEMCELL technologies at room temperature for 20 min. About 100 – 150 isolated crypts per well were seeded in Matrigel with ENR (Egf, Noggin and R-Spondin1) or WENR (Wnt3a-containing ENR) + Nicotinamide (WENR+Nic) culture medium composed of advanced Dulbecco’s modified Eagle medium (DMEM)/F12 supplemented with penicillin/streptomycin, 10 mM HEPES (Gibco), GlutaMAX (Gibco), 1x B27 (Life Technologies), 10 mM nicotinamide (MilliporeSigma; used only in WENR), 1.25 mM N-acetylcysteine (Sigma-Aldrich), 50 ng/ml mEGF (Peprotech), 100 ng/ml mNoggin (Peprotech), 10% R-spondin1 conditioned medium, 50% Wnt3A conditioned medium (only in WENR), and 10 mM ROCK inhibitor (Tocris). Established organoids were routinely passaged at a 1:3 – 1:5 ratio every week and maintained in culture medium without ROCK inhibitor.

### Organoid single-cell analysis

To induce gene recombination in Vil-CreERT2 Daam1/2 double conditional knockout (D1/2 cDKO) and Vil-CreERT2 Rnf43-Znrf3 double conditional knockout (RZ cDKO) small intestinal organoid lines, 1 μg/ml 4-hydroxytamoxifen (4-OHT) was added overnight in WENR+Nic (Wnt, Egf, Noggin, R-spondin1 + nicotinamide) organoid culture medium. After recombination, D1/2 and RZ mutant organoids were cultured in WEN+Nic medium for selection purposes, as only successfully recombined mutant cells survive in the absence of Rspo1. WT, D1/2 cDKO and RZ cDKO organoids were maintained in regular WENR+Nic (WT) or WEN+Nic (D1/2 and RZ cDKO) culture medium for 10 days prior to single-cell analysis. Three days prior to single-cell analysis, organoids were passaged at a 1:3 ratio. For analysis, organoids were mechanically and chemically dissociated and prepared for library generation and sequencing by Vienna BioCenter Next Generation Sequencing Facility. WT (8,000), RZ cDKO (13,000), and D1/2 cDKO (9,000) cells were analyzed and obtained sequencing data were processed as described below.

### scRNA-seq data preprocessing

To generate count matrices for each organoid sample such as wildtype (‘WT_WENR’), Rnf43/Znrf3 double knock-out (RZ_WEN), and Daam1/Daam2 double knock-out (Daam_WEN), Cellranger (v6.1.1)^37^ was utilized with default option and mouse reference (‘refdata-gex-mm10-2020-A’) provided in 10X genomics. Based on Seurat pipeline (v4.1.0)^38^, we generated analytic objects from each gene-by-cell matrix from ‘filtered_feature_bc_matrix’ of cellranger, we firstly filtered-out poor-quality cells with less than 4,000 and more than 40,000 unique molecular identifiers (UMIs), more than 13% percent mitochondrial genes, and more than 4,000 detected genes for WT_WENR and RZ_WEN samples. Especially for Daam_WEN sample, we applied partially different criteria to discard poor-quality cells with more than 50,000 UMIs and more than 5,000 detected genes. After quality control step, we had 3,828 cells (WT_WENR), 4,342 cells (RZ_WEN), and 3,257 cells (Daam_WEN). Then, we sequentially conducted NormalizeData (log-normalization) and FindVariableFeatures (‘vst’ method, 2,000 features) functions of Seurat pipeline for each sample. In order to select singlet, DoubletFinder_v3, which is a recent version of DoubletFinder^39^, was conducted with default parameters. Using cell cycle markers derived from AnnotationHub R package (v3.4.0) with ‘Mus musculus’, ‘EnsDb’ for S and G2/M phases, we calculated cell cycle scores via CellCycleScoring function and regressed out based on the scores such as ‘S.Score’ and ‘G2M.Score’ on Seurat pipeline. After merging three count objects and storing each object to sample category, we performed PCA with RunPCA function to prepare harmonization which is one of batch reduction methods. The merged object was harmonized through RunHarmony function with ‘Sample’ as biological factor and 30 PCs^40^. Based on the dimensionally reduced object, we identified clusters through the sequential procedures such as RunUMAP, FindNeighbors (‘k.param’ is 25), and FindClusters (‘resolution’ is 0.75, Louvain algorithm) with 30 harmony PCs.

### scRNA-seq data analysis

In order to annotate cell types in the dataset, we used cell type markers described in Extended Data Fig. 5. Using expression profile of clusters, the clusters have been combined to reflect that intestinal organoids cultured in conventional media consist of a large proportion of stem cells and a small proportion of fully differentiated cells^41^. Because genetic change may affect to cell homeostasis and transcriptomic changes of cell type markers, we annotated cell types based on gene expression in WT_WENR. In order to generate dot plots for each sample origin, expression patterns of each gene within cells corresponding to sample and cluster instance were used. Using dittoBarPlot function in dittoSeq R package (v1.6.0)^42^, we generated cell type proportion in each sample. The statistical significances for the difference of a cell type between samples were estimated through Fisher’s exact test. We showed expression pattern of each gene on UMAP clusters for combined samples and individual samples based on UMAP coordinates and expression level.

### Pseudo-bulk RNA-seq analysis

From single cell RNA-seq data, aggregation of UMI counts from group of cells has been used to observe differential gene expressions across clusters or sample-wise. In order to capture global effect of genetic differences in transcriptome, we adopted pseudo-bulk approach with 10 pseudo-samples from each of scRNA-seq count matrices with QC passed cells according to WT_WENR, RZ_WEN, and Daam_WEN samples. Each sample has a gene-by-pseudo-sample count matrix and each column (i.e., a pseudo-sample) contains the aggregated UMI counts for the corresponding genes from same number of randomly selected cells. We employed DESeq2^43^ R package to normalize UMI counts. For each gene, we compared log-scaled normalized expression levels from samples and estimated statistical significance with Mann-Whitney U test and Kruskal-Wallis test by using ‘stat_compar_means’ function on ‘ggpubr’ R package (v0.4.0).

### Organoid electroporation

Small intestinal organoids were electroporated following a previously established protocol^15^. Two days before electroporation, culture medium was replaced from WENR+Nic to EN+Nic in the presence of ROCK inhibitor and the Gsk3 inhibitor CHIRON 99021. One day before electroporation, 1.25% v/v DMSO was added to the culture medium. On the day of electroporation, organoids were dissociated into small clusters by TrypLE treatment and electroporation was performed in 400 μl of BTXpress buffer (Havard Apparatus) with 12.5 μg of DNA using a NEPA21 Electroporator (NEPA Gene). For CRISPR/Cas9 mediated knockout, organoids were electroporated with CRISPR-concatamer vectors containing gene-specific guide RNAs (gRNAs) in combination with a Cas9 expression plasmid (Addgene #41815), at a 1:1 ratio. The electroporated organoids were seeded in Matrigel and cultured with EN medium in the presence of Nic and ROCK inhibitor. Two days later, post-electroporation culture medium was replaced by regular culture medium (ENR or WENR). DNA oligos and primers used to generate and analyze CRISPR KO organoids are listed in Table 1.

**Table 1.**
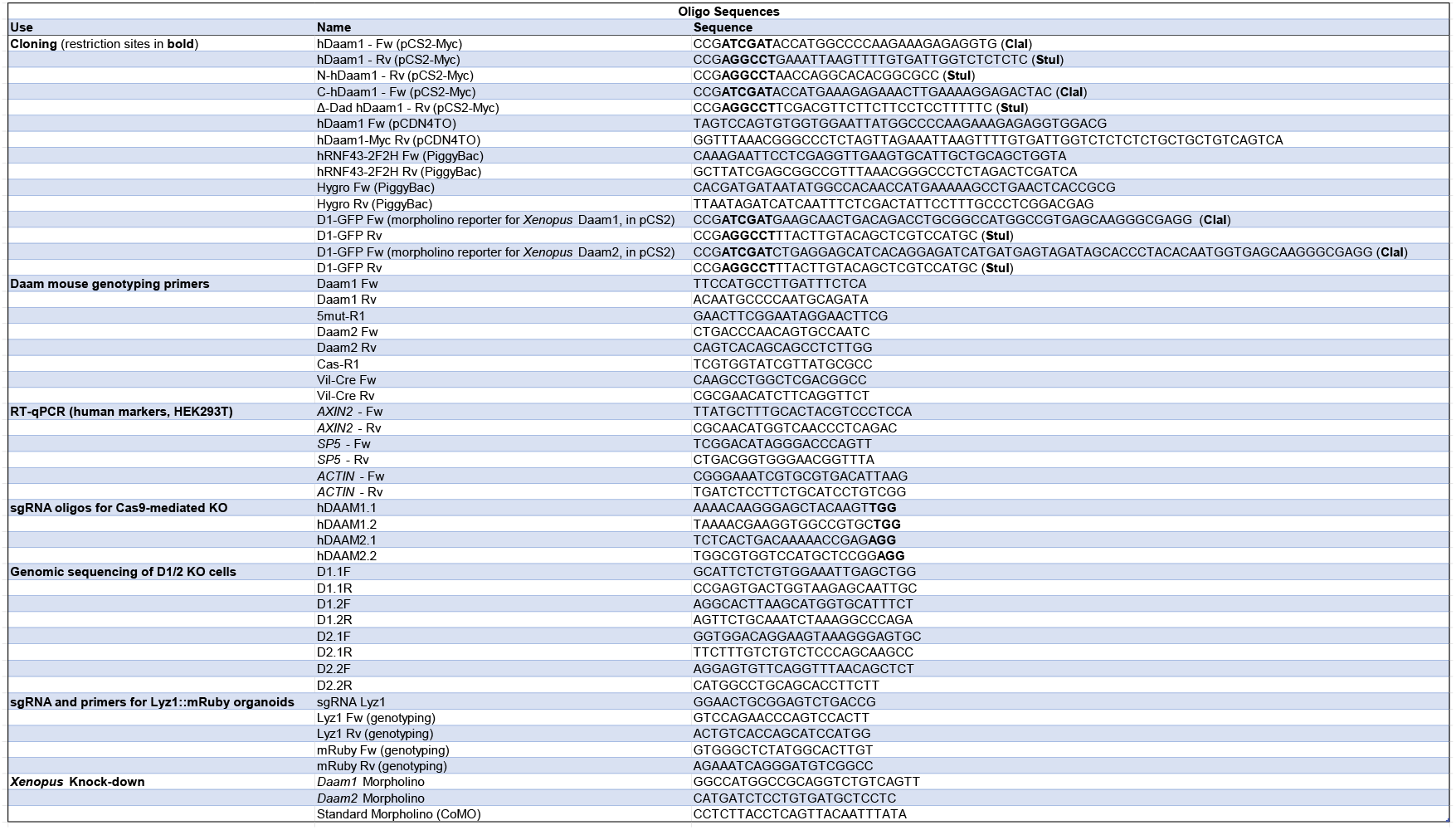
List of DNA oligos and primers used in this study.

### Generation of a Paneth cell reporter SI organoid line

SI organoids expressing tamoxifen-inducible Cas9 were established from Vil-CreERT2; Rosa26 - floxed STOP - Cas9 mice. These organoids were co-electroporated with a plasmid containing *Lysozyme1* (*Lyz1*) sgRNA for *Lyz1* and a Golden Gate-generated targeting vector containing the coding sequence of *2A-peptide-mRuby* red fluorescent protein, followed by a PGK-driven blasticidin resistance gene and flanked by homology arms (HA) of 48 bp of length, that are homologous to the region immediately up- and downstream of the Lyz1 stop codon. After electroporation, organoids were selected with 100 μg/ml blasticidin for 6-7 days. Hence, single surviving organoids were manually isolated to generate monoclonal lines and genotyped to confirm the correct integration of *mRuby* in *Lyz1*. For Wnt5a treatment, Lyz1::Ruby organoids were seeded in ENR, WENR or W5aENR, where recombinant Wnt5a protein (rWnt5a, R&D Systems) is added to ENR at a final concentration of 200 ng/ml. Organoids were cultured for 5 days before imaging analysis. Media were refreshed every 2 days.

### Immunohistochemistry and RNA-Scope *in situ* hybridization

All immunohistochemical (IHC) staining experiments on mouse intestinal sections were performed by the IMBA Histology Facility at Vienna BioCenter Core Facilities (VBCF), member of the Vienna BioCenter (VBC), Austria. All samples were incubated in 3% H_2_O_2_ in blocking solution [2% BSA, 5% goat serum, 0.3% Triton-X100 in PBS (phosphate buffered saline, 137 mM NaCl, 2.7 mM KCl, 10 mM Na_2_HPO_4_, 1.8 mM KH_2_PO_4_, pH 7.4)] at room temperature (RT) for 10 min and further incubated in blocking solution at RT for 1 hour. Sections were incubated overnight with primary antibodies diluted in blocking solution, followed by 3 washes in PBS and incubation with secondary antibody solution for 1 hour at RT. After 3 additional washes in PBS, sections were processed for Hematoxylin/Eosin (H&E) staining, which was performed without heat using the Epredia Gemini AS Automated Slide Stainer, and finally mounted. For RNAScope/IHC protocol, 4 μm thick sections were processed using the RNAscope® Multiplex Fluorescent Detection Kit (ACDBio), following manufacturer’s directions. Finally, stained slides were mounted with fluorescence mounting medium (Dako).

### Generation of DAAM1 and PiggyBac RNF43 expression vectors

Human *DAAM1* cDNA was purchased from TransOMIC. Full length or truncated *DAAM1* constructs were PCR-amplified using Phusion High Fidelity Polymerase (NEB) and cloned into pCDNA4/TO or pCS2 expression vectors, containing a Myc Tag for C-terminal fusion. In order to generate pBhCMV-hRNF43-IRES-mCherry and pPB-CAG-rtTA_Hyg, cDNAs of human RNF43 and hygromycin resistance were PCR-amplified as mentioned above and cloned into PiggyBac-based vectors containing tet-responsive elements, using the In-Fusion HD cloning kit (Clontech), according to manufacturer’s instructions. RNF43 and rtTA expression constructs were electroporated into SI organoids as described above, always in combination with the Super PiggyBac Transposase expression vector in a 2:2:1 ratio. All constructs reported here were sequence-verified using Sanger sequencing.

### Cell culture, DNA transfection and growth factor stimulation

HEK293T cells were maintained in DMEM supplemented with 10% FBS, 1% glutamine and 1% penicillin-streptomycin, kept in a 37 °C and 5% CO_2_ incubator and passaged every 5 – 7 days. Transfection was performed by using 1 μg/ml polyethylene imine (PEI), pH 7.4, and plasmid DNA at a 5:1 ratio^44^. Plasmid DNA used was 5 μg per 6 cm culture dish (ubiquitination and Fzd degradation assays) or 10 μg per 10 cm dish (co-immunoprecipitation experiments). For Fzd degradation and ubiquitination analysis, Rnf43 was transfected at a ratio of 1:5 to total plasmid DNA. For Wnt3a treatment (as shown in Fig. 4c and d), HEK293T cells were plated on 12-well plates, and upon reaching 80% confluency cells were treated with Wnt3a-conditioned medium (CM), overnight for real time-quantitative PCR or for the time indicated in the case of active β-catenin immunoblot. For the experiment shown in Extended Data Fig. 3a, endogenous Wnt ligand secretion was inhibited by overnight treatment with 1 μM LGK-974 (PeproTech). Subsequently, the non-canonical Wnt pathway was activated by adding rWnt5A (R&D Systems) to the culture medium at 40 or 80 ng/ml final concentration, with or without 50 ng/ml of recombinant human R-Spondin-1 (PeproTech). For Wnt5a-induced actin cytoskeleton rearrangements, untransfected HEK293T cells or cells transfected 48 hours earlier with pEGFP-C1 Lifeact-EGFP (Addgene, plasmid #58470) were treated with Wnt5a CM or protein (200 ng/ml) for 2 hours at 37 °C, before fixation and immunostaining. Wnt5a CM was produced from commercially available L-cells (ATCC, CRL-2814), while Wnt3a CM was produced from L-cells kindly provided by Hans Clevers (Hubrecht Institute, Utrecht, Netherlands), following standard protocols. WT L-cells (ATCC, CRL-2648) were instead used to produce control CM.

### HEK293T Daam1/2 DKO clone generation

To generate Daam1/2 DKO clones, CRISPR/Cas9 mediated knockout was performed as follows. WT HEK293T cells were seeded 3 days before transfection and a concatemer construct harboring Daam1 and Daam2 sgRNAs and Cas9-expressing vector were co-transfected, together with GFP. Successfully transfected HEK293Tcells were sorted by FACS and seeded as single cells to confirm genotype of Daam1 and Daam2 by Sanger sequencing followed by TIDE analysis for the quantitative assessment of CRISPR gene editing. DNA oligos and primers used are listed in Table 1.

### Immunofluorescence and confocal imaging

Whole mount immunostaining on organoids was performed following the protocol described by van Ineveld et al.^45^, with no modifications. For HEK293T immunofluorescence, cells were grown on 12-well plates containing glass coverslips, pre-coated with a solution containing 0.01% poly-L-Ornithine (Millipore) overnight at 37°C. Cells were then transfected and/or stimulated as described above, followed by two washes with PBS, fixation in 4% (wt/vol) paraformaldehyde (PFA) in PBS for 20 min, then permeabilization with 0.2% (vol/vol) Triton X-100 in PBS. Coverslips were then washed with PBS, blocked for 1 hour in blocking buffer consisting of 3% (wt/vol) BSA in PBS at room temperature, and incubated with primary antibodies in blocking buffer overnight at 4°C. The next day cells were washed 3 times with PBS, incubated with secondary antibodies diluted in blocking buffer for 1 hour at room temperature and mounted onto glass slides with ProLong Gold antifade reagent with DAPI (Life Technologies) to stain cell nuclei. A list of primary and secondary antibodies used in this study, including their dilution, is provided in Table 2. Cells and organoids were imaged using an inverted LSM 880 Airyscan confocal microscope (Carl Zeiss, Jena, Germany) using 405-, 488- and 561-nm lasers for excitation, and a 20× objective (Plan Apochromat × 20/0.8). Z-stacks were acquired with a resolution of 1024 pixels, snaps with resolution of 2048 (frame size 2048×2048). For scanning, the following parameters were used: unidirectional scanning, averaging number 8, 8 bit depth. Images were acquired with multi-tracking for each fluorophore and Zeiss ZEN Black Edition software. ZEN Blue Edition software was used for image analysis. The list of antibodies and fluorophores used for immunostaining throughout this paper is provided in Table 2.

**Table 2.** List of antibodies or fluorescent labels used for immunofluorescence in this study.

| Immunofluorescence |  |  |  |
| --- | --- | --- | --- |
| Primary Antibody | Company | Catalog Number | Dilution |
| Mouse anti-Daam1 (WW-3) | Santa Cruz Biotechnology | sc-100942 | 1:250 |
| Mouse anti-Daam2 (E-1) | Santa Cruz Biotechnology | sc-515129 | 1:250 |
| Mouse anti-Ki67 (B56) | Abcam | ab279653 | 1:300 |
| Rabbit anti-human Lysozyme EC 3.2.1.17 | Dako | A009902-2 | 1:500 |
| Secondary Antibody or Fluorescent Label |  |  |  |
| Phalloidin Atto 488 | Sigma | 49409-10NMOL | 1:50 |
| SNAP-Surface 549 | NEB | S9112S | 1:1000 (1 μM final) |
| Ulex Europaeus I Agglutinin, FITC conjugated | Szabo Scandic | VECFL-1061/2 | 1:500 |
| Goat anti-mouse IgG Alexa Fluor Plus 488 | ThermoFisher | A32723 | 1:500 |
| Donkey anti-mouse IgG Alexa Fluor 546 | ThermoFisher | A10036 | 1:500 |
| Donkey anti-rabbit IgG Alexa Fluor 568 | ThermoFisher | A10042 | 1:500 |

### Surface Fzd5 internalization assay

SNAP-tagged Frizzled5 (SNAP-Fzd5) subcellular localization was monitored in wild type and Daam1/2 KO HEK293T cells in the presence and absence of Rnf43 co-expression, as previously described^6^. SNAP-surface-Alexa549 (NEB) was applied to label surface SNAP-Fzd5 for 15 min at RT in the dark, following manufacturer’s instructions. The labelled Fzd5 was chased for 30 min, after which, cells were fixed and processed for confocal imaging as described above.

### Cell surface biotin labeling

Cell surface biotinylation was performed as previously described^26^. Briefly, HEK293T cells were grown on 6-well plates previously coated overnight with a 0.1 mg/ml solution of poly-L-ornithine (Millipore), an important requirement to prevent cell detachment. 48 hours after transfection with the DNA constructs indicated in Fig. 2c, cells were transferred on ice and washed twice with ice-cold phosphate-buffered saline (PBS). Cell surface proteins were then biotinylated by incubation with 1 mg/ml of the cell membrane-impermeable reagent EZ-Link Sulfo-NHS-SS-Biotin (ThermoFisher) dissolved in PBS for 30 minutes at 4 °C with gentle agitation. Cells were washed three times with ice-cold quenching solution (50 mM glycine in PBS, pH 7.4), and twice with ice-cold PBS. Cells were kept on ice for the entire length of the labeling protocol. After the final wash, cells were lysed directly on the plates with 300 μl TNE lysis buffer (Tris-NaCl-EDTA, 50 mM Tris HCl, pH 7.4, 150 mM NaCl, 1 mM EDTA, 1% Nonidet P-40) supplemented with protease inhibitors (Roche). Cell lysates were spun at 13,000 rpm at 4 °C on a table-top centrifuge to remove cell debris, and then incubated with streptavidin-agarose magnetic beads (ThermoFisher) for 3 h at 4 °C using a head-over-head rotator to bind biotinylated proteins. An aliquot of the original cell lysate was saved for input control. At the end of the pull-down, beads were extensively washed with TNE buffer at 4 °C with rotation, and then heated to 95 °C in the presence of 2x Laemmli buffer (BioRad) to elute proteins, followed by SDS-PAGE/Western blot analysis. To assess pull-down specificity, a no-biotin control sample was included, and endogenous transferrin receptor (TfR) was monitored as a negative control.

### Immunoprecipitation

To assess Rnf43 and Daam1 interaction, we performed immunoprecipitation (IP) assays as previously described^46^, with some modifications. Briefly, HEK293T cells seeded onto 10-cm plates were transfected with the constructs indicated in Fig. 2d-f, using PEI and 10 μg total DNA. 48 hours after transfection, cells were lysed in 1 ml IP lysis buffer (10 mM Tris-HCl, pH 7.5, 100 mM NaCl, 2 mM EDTA, 1 mM EGTA, 0.5% (v/v) NP-40, 10% (v/v) glycerol) supplemented with protease inhibitor cocktail (cOmplete, EDTA-free, Roche). Cell lysates were clarified by centrifugation (13,000 rpm, 4 °C) and then pre-cleared with 20 μl of protein A/G PLUS agarose beads (sc-2003, Santa Cruz Biotechnology) for 1 hour at 4 °C, on a head-over-head rotator. Beads were removed by centrifugation and cleared lysates were then incubated with 2 μg of primary antibody (anti-HA, ThermoFisher) overnight at 4°C on a head-over-head rotator. Supernatants were then incubated with 40 μl of protein A/G agarose beads for 2 hours at 4 °C on rotation, extensively washed in lysis buffer, resuspended in 40 μl of SDS 2x Laemmlibuffer (BioRad) and heated for 5 minutes at 95 °C to elute immunocomplexes, followed by analysis through SDS PAGE and Western Blot. For IP/mass spectrometry (IP/MS) to identify Rnf43 interactors, Rnf43-2xFlag-2xHA was pulled-down with anti-Flag agarose beads (Sigma). A total of 5 independent experiments were included in this assay, and a Rnf43 deletion construct containing only the extracellular protease-associated and transmembrane domains (PA/TM Rnf43) was used to assess for interaction specificity. MS on immunoisolated complexes was conducted as previously described^6^. The hit list containing all identified peptides and their relative abundance is provided in Supplementary Table 1.

### Ubiquitination assay

HEK293T cells seeded in 6-cm plates and maintained in DMEM with 10% FBS and penicillin-streptomycin were transfected with ubiquitin-Myc-6xHis, V5-Frizzled5 and Rnf43-2xFlag-2xHA using PEI. Where required, transfected cells were incubated with 10 nM Bafilomycin A1 (BafA1) overnight before lysis. Two days after transfection, samples were harvested and lysed with 500 μl IP lysis buffer (as described above) supplemented with protease inhibitor cocktail (cOmplete, EDTA-free, Roche) and 10 mM N-Ethylmaleimide (NEM, Sigma-Aldrich, E3876). Lysates were then incubated with V5-Trap magnetic agarose beads (ChromoTek) overnight at 4 °C before being processed for Western Blot analysis.

### Wnt5a-induced active Rho assay

Active Rho pull-down was performed as previously described^47^, with some modifications. HEK293T cells were stimulated with rWnt5a at a concentration of 200 or 400 ng/ml for 30 minutes at 37 °C. Next, cells were quickly washed with ice-cold PBS and lysed in ice-cold lysis buffer (25 mM Tris•HCl, pH 7.2, 150 mM NaCl, 5 mM MgCl_2_, 1% NP-40 and 5% glycerol). Crude lysates were then clarified by centrifugation and the supernatant was incubated with glutathione magnetic beads (ThermoFisher), conjugated with the Rhotekin Rho Binding domain (RBD)-GST fusion protein (produced from Addgene plasmid #15247), for 1 hour at 4 °C, before proceeding to Western blot analysis.

### Western Blotting

SDS-PAGE and Western blots were performed using pre-cast gradient gels (ThermoFisher), using standard protocols. Blotted nitrocellulose membranes were analyzed using the Li-Cor software Odyssey 3.0. All primary and secondary antibodies were diluted either in Tris-buffered saline plus 0.1% Tween 20 (TBST) containing 2.5% (wt/vol) of Blotting Grade Blocker (Bio-Rad), or in LiCor Intercept (TBS) blocking buffer supplemented with 0.1% Tween 20. The list of antibodies used and their dilution is provided in Table 3.

**Table 3.** List of antibodies used for Western blot. Antibodies also used for IP are in bold.

| Western Blot |  |  |  |
| --- | --- | --- | --- |
| Primary Antibody | Company | Catalog Number | Dilution |
| Chicken anti-GFP | Aves Labs | 1020 | 1:1000 |
| **Mouse anti-HA (2-2.2.14)** | ThermoFisher | 26183 | 1:1000 |
| Rabbit anti-Myc (71D10) | Cell Signaling Technology | 2278 | 1:1000 |
| Rabbit anti-V5 (D3H8Q) | Cell Signaling Technology | 13202 | 1:1000 |
| Rabbit anti-active (non-phosphorylated) β-Catenin (D13A1) | Cell Signaling Technology | 8814 | 1:1000 |
| Rabbit anti-CD71 (D7G9X) (Transferrin Receptor, TFR) | Cell Signaling Technology | 13113 | 1:1000 |
| Mouse anti-Daam1 (WW-3) | Santa Cruz Biotechnology | sc-100942 | 1:500 |
| Rabbit anti-Dvl2 (30D2) | Cell Signaling Technology | 3224 | 1:1000 |
| Rabbit anti-Dvl3 | Cell Signaling Technology | 3218 | 1:1000 |
| Rabbit anti-Lrp6 (C5C7) | Cell Signaling Technology | 2560 | 1:1000 |
| Mouse anti-ROR2 (H-1) | Santa Cruz Biotechnology | sc-374174 | 1:1000 |
| Rabbit β-actin (13E5) | Cell Signaling Technology | 4970 | 1:3000 |
| Rabbit anti-Gapdh (14C10) | Cell Signaling Technology | 85925 | 1:3000 |
| Mouse anti-α-tubulin (DM1A) | Cell Signaling Technology | 3873 | 1:3000 |
| **Secondary Antibody** |  |  |  |
| Donkey anti-rabbit IRDye 680 RD | Li-COR | 926-68073 | 1:4000 |
| Donkey anti-rabbit IRDye 800 CW | Li-COR | 926-32213 | 1:4000 |
| Donkey anti-mouse IRDye 680 RD | Li-COR | 926-68072 | 1:4000 |
| Donkey anti-mouse IRDye 800 CW | Li-COR | 926-32212 | 1:4000 |
| Donkey anti-chicken IRDye 800 CW | Li-COR | 926-32218 | 1:4000 |

### *Xenopus* husbandry and embryo injection

Wildtype frogs were obtained from the European *Xenopus* Resource Center (EXRC), UK and NASCO, USA, and group-housed at the Institute of Molecular Pathology (IMP) facilities. All animal handling and surgical procedures were carried out adhering to the guidelines of the Austrian Animal Care and Use Committee. *In vitro* fertilization was performed as previously described^48^. Briefly, testes were surgically removed from a male frog anesthetized in 0.03% tricaine methanesulfonate (MS222, Sigma-Aldrich), and a sperm suspension was obtained by crushing each testis in 1 ml of 1x Marc’s Modified Ringers (MMR, 0.1 M NaCl, 2.0 mM KCl, 1 mM MgSO4, 2 mM CaCl2, 5 mM HEPES, pH 7.4). Ovulation of female frogs was induced the night before the experiment by injecting 500 IU of human chorionic gonadotropin (hCG). On the day of the experiment, frogs were allowed to spontaneously lay eggs in a high-salt solution (1.2x MMR). Laid eggs were collected and fertilized with 200–300 μl of the sperm suspension. To remove the jelly coat, fertilized eggs were treated with 2% cysteine in 0.1x MMR, pH 7.8, for about 7 min at RT. Dejellied embryos were then cultured in 0.1x Marc’s modified Ringer’s solution and staged according to Nieuwkoop and Faber^49^. Translation-inhibiting Morpholino (MO) antisense oligos specific for *Xenopus daam1* and *daam2* were obtained from GeneTools. The *daam1* MO sequence was previously reported^50^, while *daam2* MO was designed *ex novo* for this study. *Xenopus* genomic sequence deposited at Xenbase was used to verify that the designed *daam* MOs target both L and S homeologs^51^. All morpholino sequences used in this study are listed in Table 1. For embryo injections, 30 ng total/embryo of *daam1/2* MOs were injected alone or in combination into the 2 dorsal blastomeres of 4-cell stage embryos. Standard Morpholino (CoMO, targeting a human beta-globin intron mutation that causes beta-thalassemia) was used as a negative control and injected at a similar concentration. For *wnt8* injection, 32 ng total/embryo of *pCSKA-xwnt8* (Addgene # 16866) plasmid DNA was injected into the 2 dorsal blastomeres of 4-cell stage embryos. After injection, embryos were collected at early tailbud stage, fixed in 4% PFA in PBS for two hours at RT and washed extensively with PBS to remove residual PFA. Images were acquired on a color camera-equipped stereomicroscope (Zeiss). To assess morpholino specificity, we cloned the 5’ coding or untranslated region (UTR) of *Xenopus daam1* and *2* genes, respectively, containing the morpholino binding sites, upstream of the *gfp* open reading frame (ORF), into pCS2+ vector. The presence of *daam* sequences did not affect GFP fluorescence. Plasmids were then linearized using NotI restriction enzyme and mRNA was transcribed *in vitro* using Sp6 mMessage mMachine kit (ThermoFisher), following the manufacturer’s instructions. 250 pg total of *d1/2-gfp* mRNAs were injected animally into 2 blastomeres of 4- or 8-cell stage embryos, together with 30 ng total of *daam1/2* or control MOs (as indicated in Extended Data Fig. 4). Finally, injected embryos were collected at the neurula stage and processed for Western blot analysis.

### RT-qPCR

RNA extraction from HEK293T cells and mouse small intestinal organoids, cDNA preparation and RT-qPCR were performed according to established protocols^52^, with minor modifications. Briefly, HEK293T cells from single wells of a 6-well plate were pelleted and total RNA was extracted using the RNeasy MiniKit (QIAGEN), following the manufacturer’s instructions. For organoid total RNA preparation, three single wells of a 48-well plate were pooled together after removing Matrigel with Cell Recovery Solution (Corning), 1 hour at 4 °C. 1 μg of total RNA was used for cDNA synthesis with Oligo(dT)_18-22_ primers by SuperScript III reverse transcriptase (ThermoFisher). 2 μl cDNA was then used for quantitative RT-PCR, which was performed on a CFX Connect Real-Time System thermal cycler (BioRad), using GoTaq qPCR master mix (Promega) and including 2 to 3 biological replicates for each marker and reference gene. Expression levels were normalized to housekeeping gene actin. Primer sequences are listed in Table 1.

---

**Extended Data Fig. 1.**
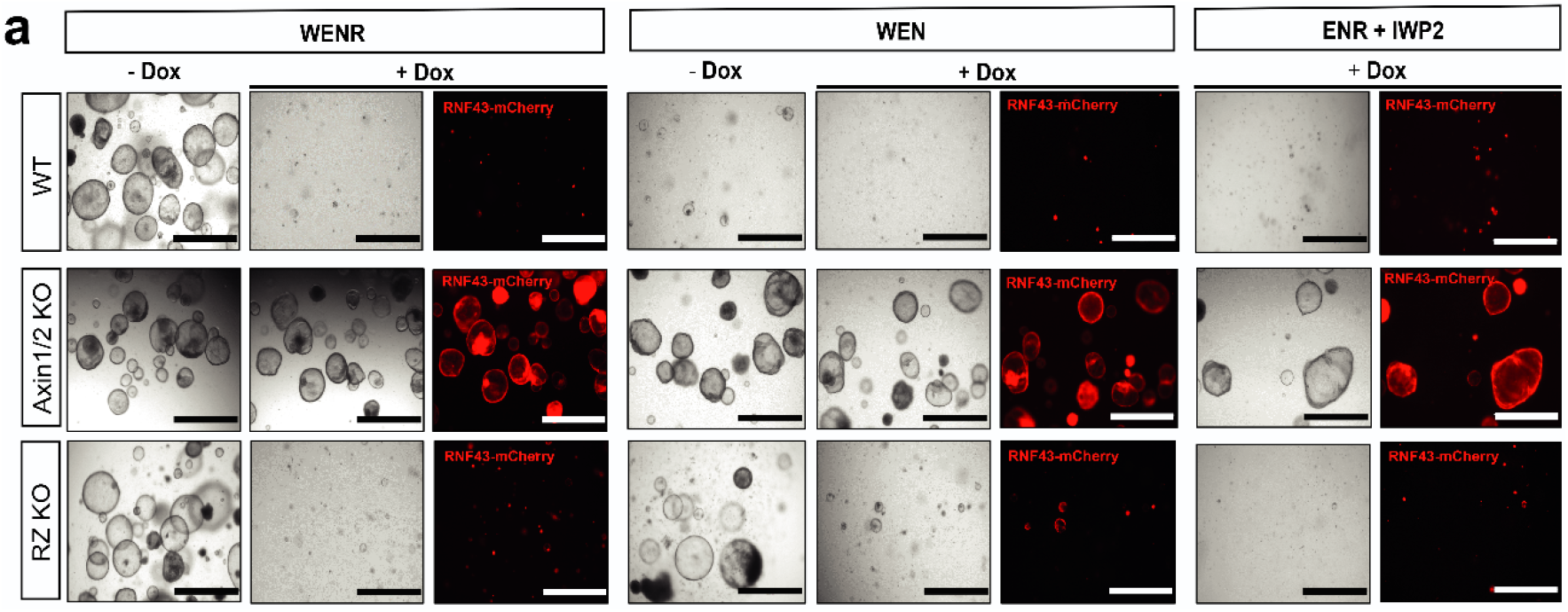
Specificity assay for *Rnf43-IRES-mCherry* overexpression in SI organoids. **a,** In contrast to WT organoids, Axin1/2 CRISPR KO mutants can survive in the absence of Rspo1 and Wnt ligands, while RZ KO organoids can only survive if Wnt is present. *Rnf43-IRES-mCherry* induction via treatment with 1 μg/ml doxycyclin causes organoids to die in all conditions, except for Axin1/2 KO due to constitutive Wnt/β-catenin signaling activation. Scale bars represent 1000 μm.

**Extended Data Fig. 2.**
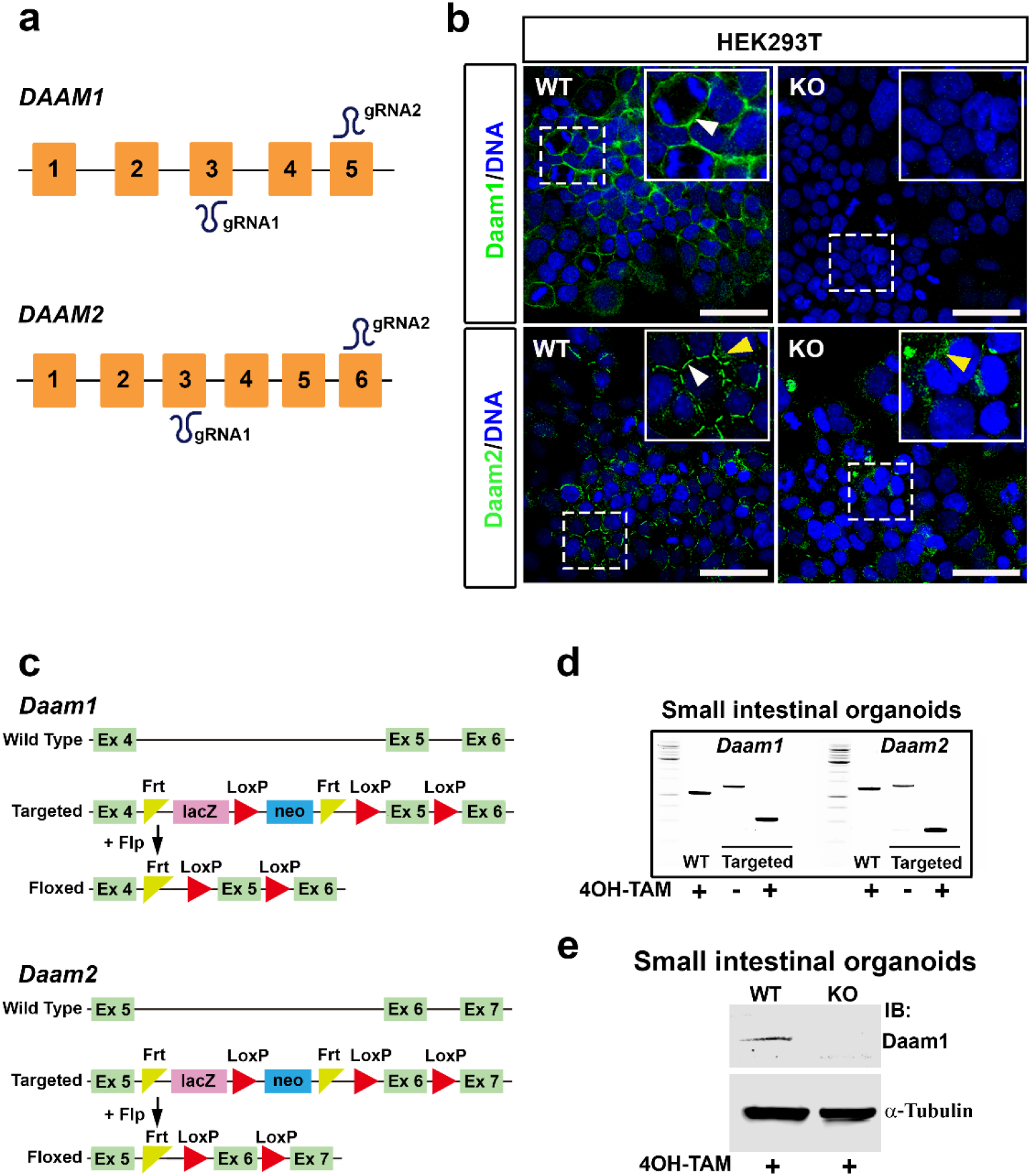
Validation of Daam1/2 knockout in HEK293T cells and mice. **a,** Diagram of the dual gRNA targeting strategy used for human *DAAM1* and *2*, showing the exons targeted by the gRNA pairs. **b,** Immunostaining for DAAM1 and DAAM2 performed in WT and D1/2 KO cells. White dashed boxed areas are enlarged in the inset, top right corners. White arrowheads point at DAAM-specific membrane staining. Yellow arrowheads indicate non-specific staining visible only with anti-DAAM2 antibody, present in both WT and KO cells. Scale bars represent 50 μm. **c,** Schematic diagram showing the targeting strategy used to generate floxed mouse *Daam1* and *Daam2* alleles, for conditional knockout. Exon 5 and Exon 6 were targeted in *Daam1* and *2*, respectively. **d,** PCR and agarose gel electrophoresis used to verify genotype and Cre-mediated recombination in D1/2 cDKO mouse organoids. **e,** Western blot confirming the absence of Daam1 protein in Vil-CreERT2 D1/2 cDKO organoids, after overnight treatment with 1 μg/ml 4-hydroxytamoxifen (4OH-TAM). After CreER induction with 4OH-TAM, organoids were cultured for at least one passage (5 days) prior to Western blot analysis. α-Tubulin was used as a loading control.

**Extended Data Fig. 3.**
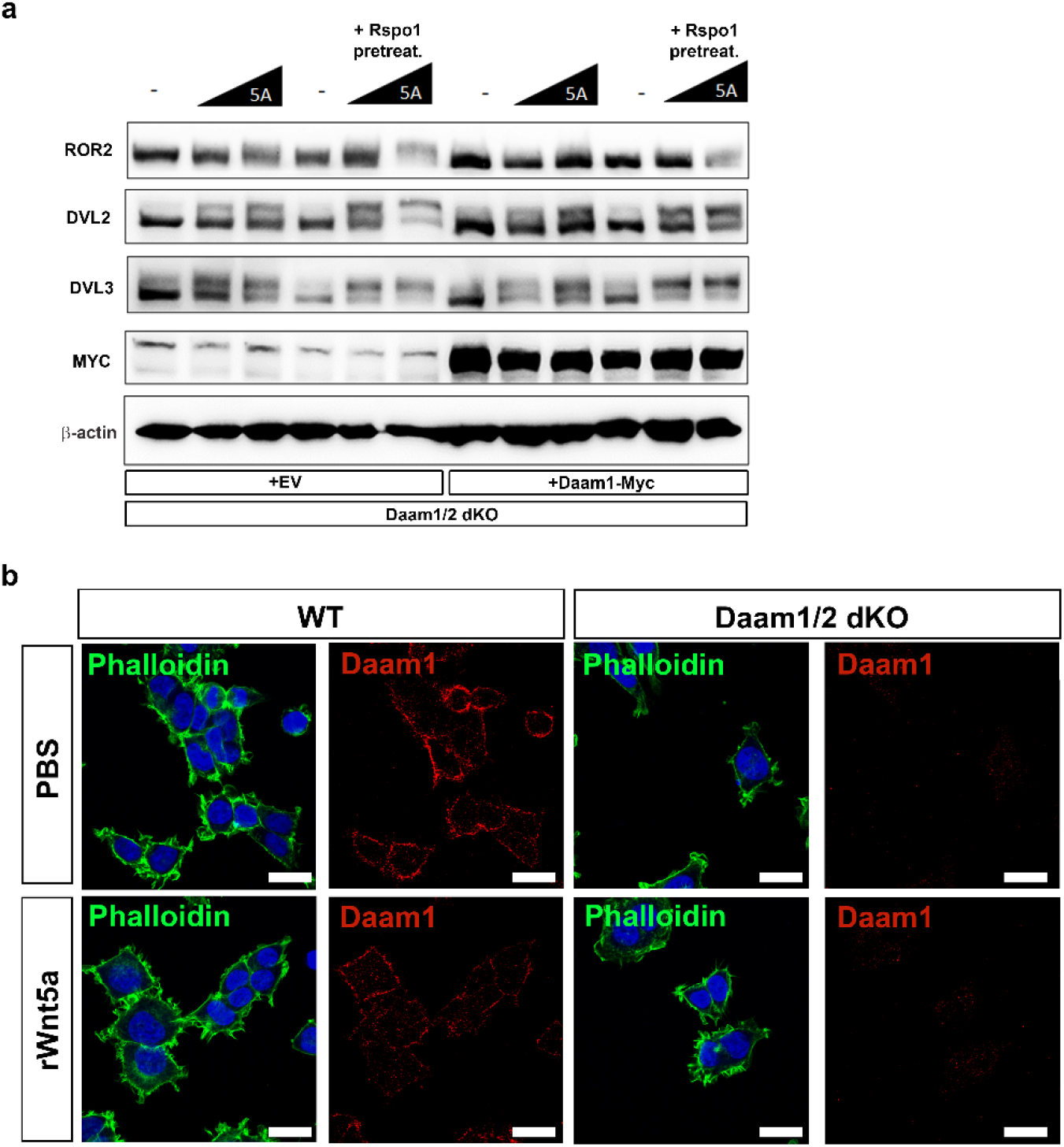
Daam1/2 KO impairs downstream events of Wnt/PCP signaling. **a,** Western blot analysis showing Wnt5a-dependent phosphorylation of ROR2 (a non-canonical Wnt co-receptor), Dvl2 and Dvl3, visualized as changes in electrophoretic mobility. Note that Wnt5a induces similar mobility shifts in both D1/2 KO cells and D1/2 KO cells transfected with Daam1-Myc. Rspo1 treatment was used to boost Wnt5a treatment, with no noticeable differences between KO and rescued samples. β-actin was used as a loading control. **b,** Confocal imaging of WT and D1/2 KO cells treated with recombinant Wnt5a (rWnt5a) protein or with PBS (mock controls), and stained for F-actin (phalloidin) and Daam1. Phalloidin reveals that actin cytoskeleton rearrangements and filopodia formation occur in WT cells upon rWnt5a treatment, but are strongly reduced in D1/2 KO cells. Daam1 protein absence in KO cells was confirmed by immunostaining. Scale bars represent 20 μm.

**Extended Data Fig. 4.**
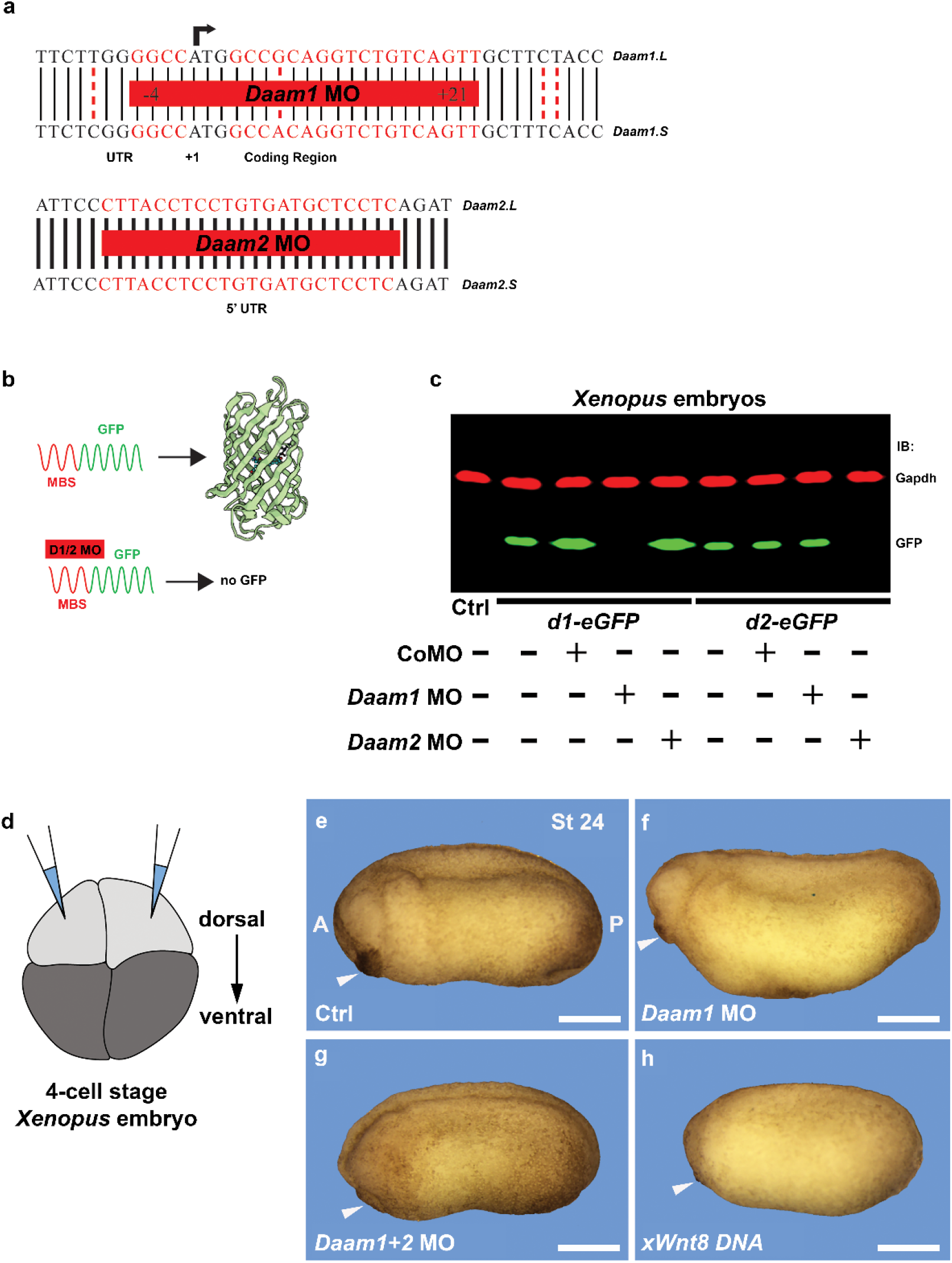
Knock-down of *Daam1* and *2* in *Xenopus laevis* embryos phenocopy canonical Wnt activation. **a,** Schematic diagram showing the *Daam1* and *2* sequences (in red) targeted by the Morpholino (MO) antisense oligos. For each *Daam* gene, MO oligos were designed to target both L and S homeolog sequences at the same time. **b,** Schematic of the experiment to assess morpholino specificity and effectiveness. MO binding sequences were cloned upstream of the GFP coding sequence, generating *d1-GFP* and *d2-GFP* constructs. **c,** Western blot of protein lysates from *Xenopus* embryos injected with *d1*- and *d2-eGFP* mRNAs. *Daam1* MO efficiently inhibits d1-eGFP translation, but not d2-eGFP, and vice versa for *Daam2* MO. Standard MO (CoMO) was used as a control. **d,** Schematic showing *Daam1/2* MO or *Wnt8* DNA microinjection into 4-cell *Xenopus* embryos. The two dorsal blastomeres of 4-cell stage embryos were targeted for injections, as these are fated to generate dorso-anterior structures, whose development is strongly regulated by Wnt activity. **e-h,** *Xenopus* embryos injected with the indicated antisense oligo or DNA plasmid and collected at stage 24 for phenotypic analysis. Ctrl embryos were injected with CoMO, which does not induce any developmental abnormality. Embryos are all oriented so that their antero-posterior (A-P) axis has the head on the left, tail on the right, as indicated in panel e. White arrowheads point to the cement gland, a prominent pigmented anterior structure, which is reduced in *Daam* morphants as well as *Wnt8*-injected embryos. Scale bars represent 500 μm.

**Extended Data Fig. 5.**
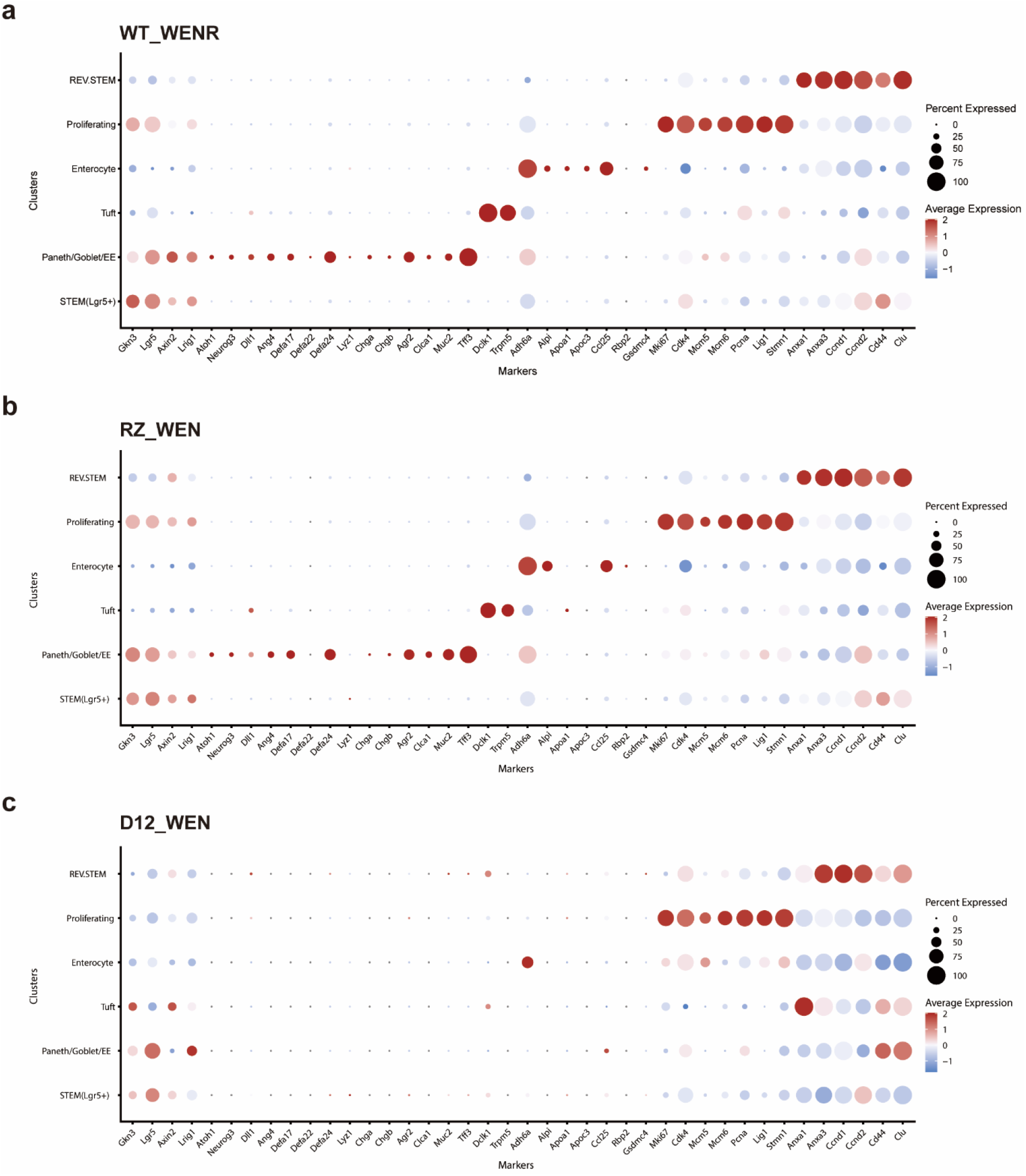
Dot plots for cell type annotation. For each cluster, the average expression levels of the corresponding gene are shown color-coded according to the scale on the right, with a gradient from blue (minimum expression) to red (maximum expression). The size of the dots indicates the fraction of cells showing expression of the corresponding genes. **a,** Wildtype organoids cultured in WENR condition. Based on the expression patterns of this sample, the integrated UMAP clusters shown in **Fig. 5** could be annotated with the different cell types. **b,** Organoids from RZ cDKO mice. **c,** Organoids from D1/2 cDKO mice.

**Extended Data Fig. 6.**
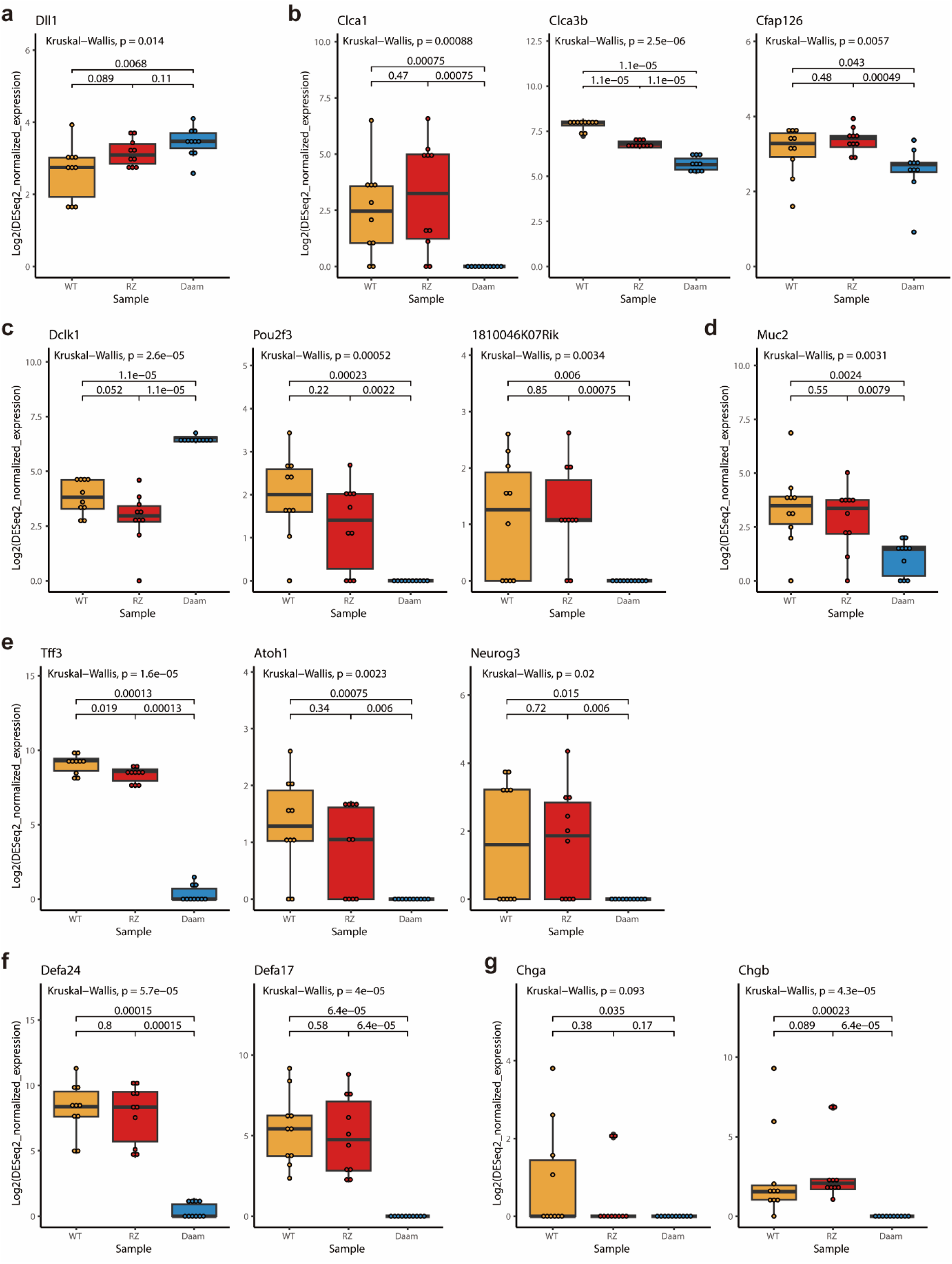
Comparisons of expression levels of cell fate marker genes with pseudobulk RNA-seq data. Normalized expression levels of corresponding genes were compared across samples. Each paired comparison was tested using the Wilcoxon test. Kruskal-Wallis test was used to test for the three samples. **a,** Dll1. **b,** Wnt/PCP target genes. **c,** Tuft cell marker genes. **d,** Goblet cell marker gene. **e,** Secretory progenitor marker genes. **f,** Paneth cell marker genes. **g,** Enteroendocrine cell marker genes.

